# Kinetoplastid-specific X2-family kinesins interact with a kinesin-like pleckstrin homology domain protein that localizes to the trypanosomal microtubule quartet

**DOI:** 10.1101/2021.09.29.462327

**Authors:** Corinna Benz, Nora Müller, Sabine Kaltenbrunner, Hana Váchová, Marie Vancová, Julius Lukeš, Vladimír Varga, Hassan Hashimi

**Affiliations:** Institute of Parasitology, Biology Center, Czech Academy of Sciences, České Budějovice, Czechia; Faculty of Science, University of South Bohemia, České Budějovice, Czechia; Laboratory of Cell Motility, Institute of Molecular Genetics of the Czech Academy of Sciences, Prague, Czechia

**Keywords:** Trypanosoma, kinesin, cytoskeleton, microtubules, microtubule quartet, morphogenesis

## Abstract

Kinesins are motor proteins found in all eukaryotic lineages that move along microtubules to mediate cellular processes such as mitosis and intracellular transport. In trypanosomatids, the kinesin superfamily has undergone a prominent expansion, resulting in one of the most diverse kinesin repertoires that includes the two kinetoplastid-restricted families X1 and X2. Here, we characterize in *Trypanosoma brucei* TbKifX2A, an orphaned X2 kinesin. TbKifX2A tightly interacts with TbPH1, a kinesin-like protein with a likely inactive motor domain, a rarely reported occurrence. Both TbKifX2A and TbPH1 localize to the microtubule quartet (MtQ), a characteristic but poorly understood cytoskeletal structure that wraps around the flagellar pocket as it extends to the cell body anterior. The proximal proteome of TbPH1 revealed two other interacting proteins, the flagellar pocket protein FP45 and intriguingly another X2 kinesin, TbKifX2C. Simultaneous ablation of TbKifX2A/TbPH1 results in the depletion of FP45 and TbKifX2C and also an expansion of the flagellar pocket, among other morphological defects. TbKifX2A is the first motor protein to be localized to the MtQ. The observation that TbKifX2C also associates with the MtQ suggests that the X2 kinesin family may have co-evolved with the MtQ, both kinetoplastid-specific traits.

## INTRODUCTION

The vermiform morphology of the unicellular parasite *Trypanosoma brucei* that causes African trypanosomiasis in humans and nagana in cattle is defined by a subpellicular array of microtubules (MTs) (Robinson *et al*., 1995; Gull, 1999; Wheeler *et al*., 2013). These MTs are cross-linked to each other and the plasma membrane via regularly spaced fibrils (Sherwin *et al*., 1987). A high order is maintained within the corset with the MT plus ends converging at the parasite’s posterior end and the minus ends located at the anterior of the cell (Robinson *et al*., 1995).

The only place where the continuity of the *T. brucei* MT array is interrupted is at the flagellar pocket, the deep invagination where the flagellum exits the cell body and the only place where endo- and exocytoses occur (Field and Carrington, 2009). The opening of the flagellar pocket is encircled by a collar, which influences the overall shape of this cavity. Distal to the collar, the so-called hook complex covers the flagellum as it exits the flagellar pocket, with two lateral arms also flanking the flagellum attachment zone (FAZ) filament and the MT quartet (MtQ) inside the cell (Albisetti *et al*., 2017). The FAZ filament and MtQ are among the interconnected fibres, filaments and junctional complexes of the multipartite FAZ (Lacomble *et al*., 2009), which together form a lateral attachment for the flagellum as it follows a helical path along the cell body, connecting the flagellar skeleton to the cytoskeleton through both the flagellar and plasma membranes (Sunter and Gull, 2016). The MtQ originates close to the basal bodies, wraps itself around the flagellar pocket and interdigitates within the MT corset next to the FAZ filament. The orientation of MTs in the MtQ is antiparallel to the one found within the MT corset, thus their minus ends are proximal to the basal bodies (Robinson *et al*., 1995).

The trypanosome MT corset is exceptionally stable and only reorganised during cell division and life cycle transitions (Sinclair and de Graffenried, 2019). Cytoskeletal structures (MtQ, MT corset, flagellum, basal bodies, which anchor the flagella within the cell, and FAZ) need to be duplicated and faithfully segregated during the cell division cycle, just like other single-copy organelles. In addition, this has to be coordinated with replication and division of the nucleus and the single mitochondrial DNA network termed the kinetoplast, which is located on the opposite side of the basal bodies relative to the flagellum (Robinson *et al*., 1995; Wheeler *et al*., 2019).

Basal body duplication and segregation are the first visible steps in the cell division cycle and essential for subsequent cytokinesis (Woodward and Gull, 1990). The new MtQ starts growing just prior to basal body maturation and once the new flagellum has grown enough to invade the flagellar pocket, morphogenetic changes result in two separate pockets enveloping both the new and old flagella (Lacomble *et al*., 2010). Once the new flagellum has been nucleated and elongated, new MTs are integrated into the array between the old and new flagella, thereby increasing the overall width of the cell (Wheeler *et al*., 2013).

With its antiparallel orientation, the MtQ together with the FAZ filament create an asymmetric seam within the MT corset, which works as a ‘cellular ruler’ and helps to determine the site of future furrow ingression during cytokinesis (Sunter and Gull, 2016). Any insults to the integrity of these structures result in cytokinesis defects and it is thus tempting to envision the MtQ as a potential specialised ‘highway’ in charge of transport and delivery of cargo to the site of important cytokinesis events. Furthermore, the endplasmic reticulum associates with parts of the MtQ, although the importance of this association remains mysterious. The first protein localized to this structure, TbSpef1, is found at the proximal end of the MtQ, which wraps around the flagellar pocket between basal bodies and the flagellar pocket collar (Gheiratmand *et al*., 2013). Its depletion causes problems with the biogenesis of the MtQ and associated structures as well as motility defects. The minus end of the MtQ MTs is anchored to the basal bodies by a protein named TbSAF1, found in close proximity to TbSpef1 but localized between the pro and mature basal bodies (Dong *et al*., 2020). TbSAF1 was found in a proximal proteomics screen of proteins adjoining to and/or directly interacting with TbSpef1, which found three additional proteins that localize to the MtQ between the basal body and flagellar pocket collar (Dong *et al*., 2020). However, no motor proteins that localise to the MtQ, and which may potentially bring cargo to the cytokinesis site, have thus far been identified.

Kinesins are well-studied motor proteins that regulate MT dynamics, organize MT networks, and transport cargo along MTs. Transporting kinesins can form homo- and heterodimers and consist of a head region containing the kinesin motor domain, which is responsible for ATP hydrolysis and MT binding, and a tail region important for dimerization with a kinesin/kinesin-like partner and cargo binding (Hirokawa and Noda, 2008). In trypanosomatid genomes, kinesins and kinesin-like proteins constitute one of the largest protein superfamilies (Berriman *et al*., 2005; El-Sayed *et al*., 2005; Ivens *et al*., 2005). To date, 47 genes encoding high-likelihood kinesins have been identified in the *T. brucei* genome (Wickstead and Gull, 2006; Wickstead *et al*., 2010b). Moreover, there is a similar number of kinesin-related proteins, which did not make the threshold due to sequence variance (Berriman *et al*., 2005). Interestingly, numerous kinesins are members of two major clades named X-1 and X-2 that are restricted to the trypanosomatid lineage (Wickstead and Gull, 2006; Wickstead *et al*., 2010b). Why the kinesin repertoire has expanded so prominently in trypanosomatids remains mysterious.

Hitherto studied kinesins of *T. brucei* have been shown to be important for various processes occurring in different cellular compartments. KinA and KinB are involved in cell cycle progression and localise to the central spindle during mitosis (Li *et al*., 2008) while TbKif13-1 localizes either to the nucleus or the mitotic spindle, where it meditates chromosome segregation during mitosis (Wickstead *et al*., 2010a). KinC and KinD are distributed throughout the cytoskeleton, where they are involved in subpellicular MT organization, basal body segregation and cytokinesis (Hu *et al*., 2012; Wei *et al*., 2013).

Some kinesins are associated with the flagellum and adjacent FAZ. KIN2A and KIN2B are kinesins involved in intraflagellar transport, a process that directly affects flagellar assembly and in turn affects cell motility and cytokinesis (Aslett *et al*., 2009; Bertiaux *et al*., 2020). The orphan kinesin KIN-E predominantly localises to the flagellar tip and is required for elongation of the new FAZ (An and Li, 2018). Another orphan kinesin named KLIF initially localises to the tip of the new, elongating FAZ and associates with the old FAZ as the cytokinetic furrow ingresses towards the posterior end, thus mediating the final stages of cytokinesis (Hilton *et al*., 2018). Other kinesins localising to specific cytoskeletal structures include FCP2/TbKinX1 and FCP4/TbKin15, which are found at the flagella connector (FC), a trypanosome-specific structure that connects the outgrowing new flagellum to the old one (Varga *et al*., 2017). Interestingly, FCP2/TbKinX1 belongs to kinesin clade X1, one of the two trypanosomatid-restricted kinesin families (Wickstead and Gull, 2006; Wickstead *et al*., 2010b).

Here we present data on the localisation and function of two members of the kinesin superfamily, TbKifX2A and TbPH1, with the former protein belonging to clade X2. Functional characterization and the unique localization of these kinesins further expands our knowledge on how these diverse and dexterous proteins affect the biology of these extremely diverse and successful parasites.

## RESULTS

### TbPH1 is a kinesin-like protein that fractionates with the *T. brucei* cytoskeleton

TbPH1 (Tb927.3.2490) is a 110 kDa protein named after a C-terminal pleckstrin homology (PH) domain, which is followed by a proximate homeodomain-like (HDL) fold and preceded by a coiled coil (CC) domain (Fig. 1A). The protein undergoes post-translational modifications, with phosphorylation of a serine at the start of the HDL (Urbaniak *et al*., 2012) and a methylarginine in the middle of the primary structure (Lott *et al*., 2013). While TbPH1 has an N-terminal kinesin motor domain, two substitutions of highly conserved amino acids within the Walker A motif (G93P and T/S95R) likely ablate the ATP hydrolase activity. Thus, TbPH1 is unlikely to be a *bona fide* kinesin motor protein.

**Fig. 1:**
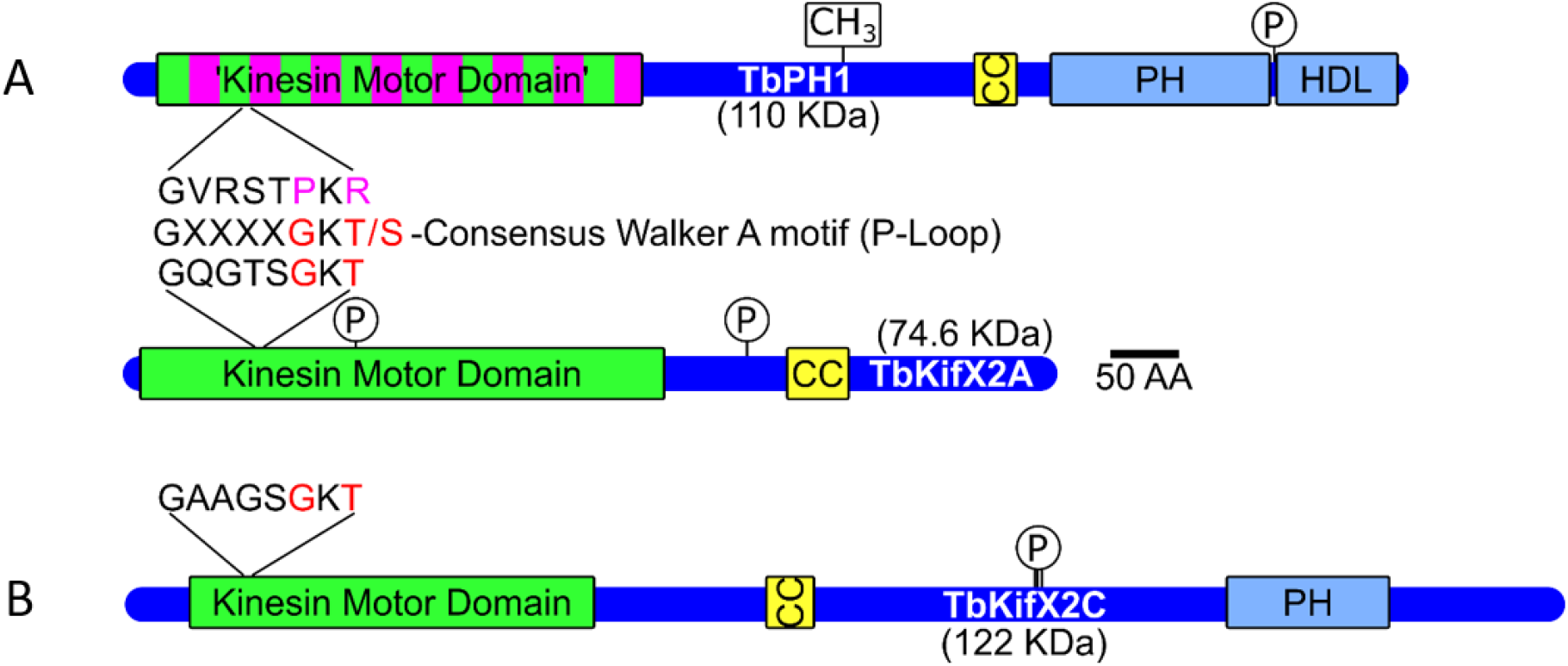
Domain architecture of TbPH1 and TbKifX2A **(A)** and TbKifX2C **(B)**. Schematic depiction includes phosphorylation and arginine methylation sites. The consensus Walker A motif is shown in between the two schemata in A for comparison of the respective motifs within the kinesin motor domains of TbPH1 and TbKifX2A. Key residues in ATP hydrolysis shown in red and mutations in purple. The likely inactive state of the TbPH1 ‘kinesin motor domain’ is highlighted by striped pattern. CC: coiled coil domain, PH: pleckstrin homology domain, HDL: homeodomain-like fold.

TbPH1 also belongs to the ∼2000 proteins found in the granule-enriched fraction (containing cellular aggregates, nuclei, kinetoplasts and flagella as well as associated cytoskeletal structures) obtained by ‘MT sieving’, a method originally conceived to purify stress granules from *T. brucei* (Fritz *et al*., 2015). In brief (see S1 Fig. for ‘MT sieving’ pipeline), cells are lysed by non-ionic detergent in low ionic strength buffer, which maintains the cytoskeleton, including the MT corset that in turn cages predominantly detergent insoluble cellular structures with at least one dimension >20-30 nm. The MT corset is then disrupted by 300 mM NaCl to release these elements for subsequent analysis.

To verify TbPH1’s apparent association with cytoskeletal fractions, we subjected a *T. brucei* cell line in which the endogenous *TbPH1* open reading frame was appended with a C-terminal V5 tag (Dean *et al*., 2015) to cell fractionation by a procedure identical to ‘MT sieving’. The cell equivalents of the ensuing fractions were inspected for the presence of TbPH1-V5 and SCD6, a stress granule marker that served as a control for this method (Fig. 2A). SCD6 was mostly found in the soluble fractions as previously reported for unstressed *T. brucei* (Fritz *et al*., 2015), with a distribution across the analysed fractions very distinct from that of TbPH1-V5. TbPH1-V5 was well retained in all cytoskeletal fractions P1 and P2, although a fraction was released in the SN1 with other soluble cytosolic proteins upon initial lysis (Fig. 2A). Importantly, an aliquot of P2 fraction that was fixed on microscopic slides for indirect immunofluorescence (IFA) using an anti-V5 antibody shows that TbPH1-V5 is localized as a line extending from the DAPI-stained kinetoplast (Fig. 2B) that decreases in signal intensity distally, suggestive of a cytoskeletal localization. This was confirmed by the observation that the vast majority of TbPH1 was released into the soluble SN3 fraction after high salt disintegration of the cytoskeleton. A small amount of TbPH1-V5 was still found in the final pellet P3, which is consistent with its detection in the cohort of proteins retained by ‘MT sieving’ (Fritz *et al*., 2015). Thus, a significant portion of the kinesin-like protein TbPH1 fractionates with the trypanosomal cytoskeleton.

**Fig. 2:**
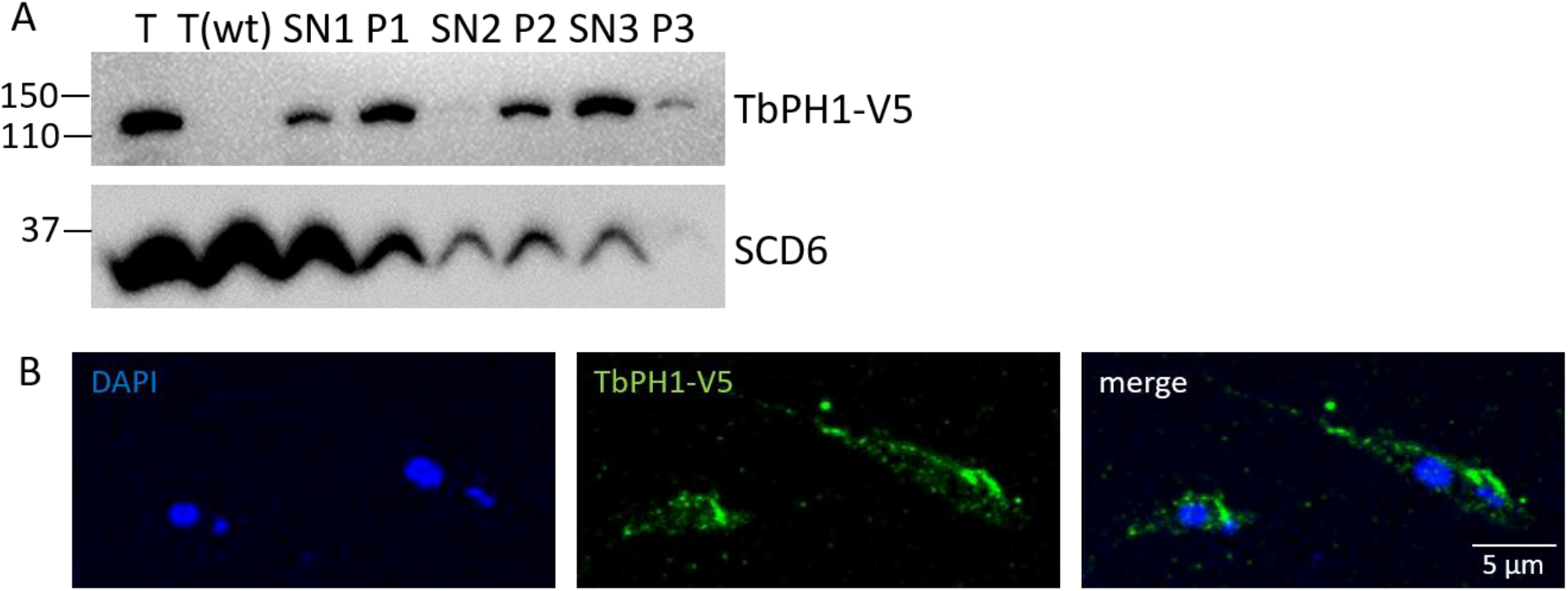
TbPH1-V5 fractionates with the cytoskeleton. **(A)** Western blot of-trypanosome cell fractions with equal cell equivalents loaded per lane as previously described in (Fritz *et al*., 2015). Fractionation procedure is summarized in S1 Fig. Antibodies used were anti-V5 and anti-SCD6 (Kramer *et al*., 2008). T: total lysate, SN: supernatant, P: pellet. **(B)** IFA of on an aliquot of fraction P2 placed on microscope slide. Cells were labelled with anti-V5 antibody (green) and DAPI (blue). The scale bar corresponds to 5 µm.

### TbPH1 strongly interacts with the trypanosomatid-restricted orphan kinesin TbKifX2A

Because kinesins tend to form dimers (Marx *et al*., 2009) and kinesin-like TbPH1 contains a CC domain (Fig. 1A), which usually mediates protein-protein interactions (Burkhard *et al*., 2001), we proceeded to see if TbPH1-V5 has an interaction partner by immunoprecipitation (IP) via its C-terminal epitope tag. To enrich for soluble TbPH1-V5 and augment IP stringency, we followed the previously described fractionation technique and used fraction SN3 as the input (Fig. 3A). A mock IP control using the parental cell line without an expressed V5 epitope was performed in parallel.

**Fig. 3:**
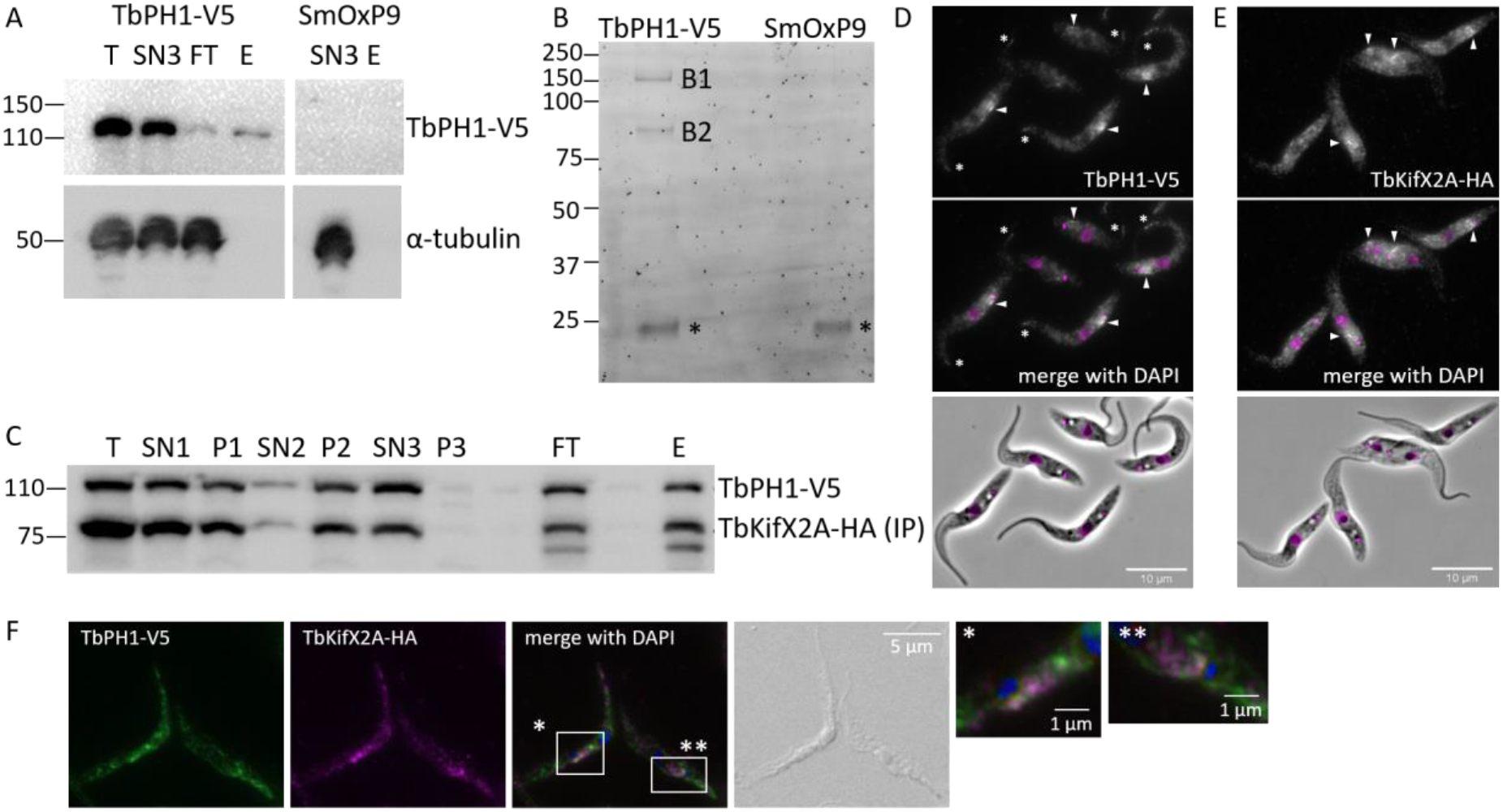
TbPH1 interacts with TbKifX2A, a kinetoplastid-specific kinesin. **(A)** Western blot verifying immunoprecipitation of TbPH1-V5. The blot was probed with anti-V5 and anti-α-tubulin antibodies. The parental cell line SmOxP9 was used as a control. T: total lysate, SN3: supernatant 3, FT: flow through, E: elution. **(B)** SyproRuby-stained gel of elution fractions of TbPH1-V5 IP and mock control (SmOxP9). B1 and B2: bands excised and by mass spectrometry identified to be TbPH1 and TbKifX2A, respectively. Asterisks: band corresponding to denatured immunoglobulin light chain that was incidentally eluted. **(C)** Western blot showing interaction of TbPH1-V5 and TbKifX2A-HA. TbKifX2A-HA was immunoprecipitated in the same way as PH1-V5. Lanes loaded as in Fig. 2A. T: total lysate, SN: supernatant, P: pellet, FT: flow through, E: elution. **(D)** IFA on whole cells detecting TbPH1-V5 (white) and counterstained with DAPI to visualize DNA (magenta). Arrowheads denote increased signal in the area along the proximal part of the flagellum. Asterisks demark an increased signal at the anterior cell body end. Merge of DAPI and Phase contrast images shown on bottom. Scale bar, 10 μm. **(E)** IFA on whole cells detecting TbKifX2A-HA (white). Images depicted as in **D. (F)** IFA on whole cells detecting TbPH1-V5 (green) and TbKifX2A (magenta). The merged image is also counterstained with DAPI (blue) to visualize DNA. Scale bar, 5 μm. Close up views of parts with strongest signals (white boxes) also shown. Scale bar, 1 μm

After confirming immunocapture of TbPH1-V5 in the first eluate (Fig. 3A), the IP eluate from this and the mock control were resolved on a SDS PAGE gel. Sypro Ruby protein gel staining showed two prominent bands of ∼117 kDa (Fig. 3B, B1), the expected size for TbPH1-V5, and of ∼80 kDa (Fig. 3B, B2). Importantly, these bands were not observed in the mock control, indicating their presence is not due to unspecific binding to the magnetic beads used to immobilize the anti-V5 antibody. Mass spectrometry analysis of these two excised bands identified TbPH1 itself as expected and a previously uncharacterised kinetoplastid-specific kinesin (Tb927.9.15470) (S1 Dataset), which we call TbKifX2A based on a previously proposed kinesin nomenclature (Wickstead *et al*., 2010b). In contrast to TbPH1, TbKifX2A’s kinesin motor domain contains a conventional Walker A motif (Fig. 1A). This motor domain is located on the N-terminus, indicating that TbKifX2A actively moves toward the plus-end of MTs (Hirokawa *et al*., 2009). It also contains a CC domain and two serine phosphorylation sites (Urbaniak *et al*., 2012), similarly to TbPH1.

Interaction between these two proteins was further verified by tagging TbKifX2A with a C-terminal HA tag in the TbPH1-V5 cell line. Indeed, TbKifX2A mirrored the fractionation pattern of TbPH1-V5 (Fig. 3C). As before, the SN3 fraction was used as an input for TbKifX2A-HA IP, which co-immunoprecipitated TbPH1-V5. Thus, we verified that TbPH1 and TbKifX2A exhibit a strong interaction that persists even in 300 mM NaCl.

Whole cells expressing TbPH1-V5 and TbKifX2A-HA were fixed and permeabilized prior to incubation with antibodies recognizing either epitope-tag (Figs. 3D and E). We observed that both proteins possess a considerable cell body signal that appears to be nucleus-excluded, with a prominent signal along the proximal region of the flagellum. This pattern is similar to the TrypTag localization data, which visualized mNeonGreen-tagged proteins in immobilized live cells (Dean *et al*., 2017). The cell body signal likely represents the cytosolic population of both proteins that was also observed in our fractionation data (Figs. 2A and 3C). It should be also noted that TbPH1-V5 only also displays an increased signal on the anterior tip of the cell body (Fig. 3D, asterisks). Co-localisation of TbPH1 and TbKifX2A in whole cells showed these two proteins to mainly associate with each other along the proximal region of the flagellum (Fig. 3F).

### TbPH1 and TbKifX2A localize to the microtubule quartet within the cytoskeleton

We next decided to address whether TbPH1 and TbKifX2A localize to or near the FAZ, as suggested by the most prominent signal obtained in whole cell IFAs. Cytoskeletons were detergent-extracted from the TbPH1-V5 and TbKifX2A-HA expressing cells to visualize any potential localization to discrete cytoskeletal structures. Indeed both proteins co-localized to a long structure running along the long-axis of the cytoskeleton adjacent to the flagellum (Figs. 4A and S2A). Furthermore, detergent extraction solubilized the cell body signal (Figs. 3D, E and F), confirming it represented the cytosolic population of TbPH1 and TbKifX2A (Figs 2A and 3C). The specificity of the antibodies recognizing V5 and HA on extracted cytoskeletons was verified by a lack of signal in the parental cell line lacking either of these epitopes (Fig. S2B).

**Fig. 4:**
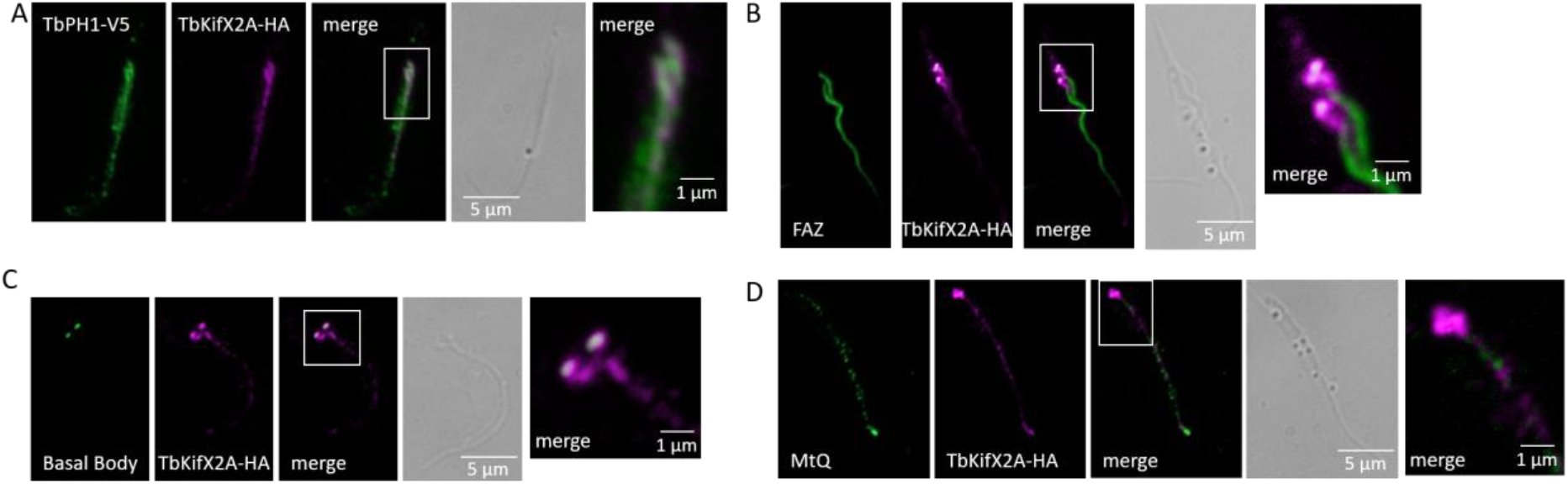
Localization of cytoskeleton-associated TbKifX2A and TbPH1. **(A)** IFA on extracted cytoskeletons probing localisation of TbPH1-V5 (green) and TbKifX2A-HA (magenta). **(B)** IFA on extracted cytoskeletons probing co-localisation of TbKifX2A-HA (magenta) with the FAZ (green), the latter labelled with L3B2 antibody. **(C)** IFA on extracted cytoskeletons probing co-localisation of TbKifX2A-HA (magenta) with basal bodies (green), the latter labelled with YL1/2 antibody **(D)** IFA on extracted cytoskeletons probing co-localisation of TbKifX2A-HA (magenta) with the MtQ (green), the latter labelled with 1B41 antibody. In all panels, merge images depict co-localization in white and DIC on right showing observed extracted cytoskeleton. Scale bar, 5 µm. Close up views of parts with strongest signals (white boxes) shown in rightmost images. Scale bar, 1 μm.

We proceeded to identify specifically which cytoskeletal element TbPH1 and TbKifX2A associate with. We used antibody recognizing TbKifX2A-HA as a proxy for the TbKifX2A-TbPH1 complex because of their demonstrated tight interaction (Fig 3), and their co-localization on detergent-extracted cytoskeletons (Fig4A, S2A). We note that in contrast to the TbPH1 co-localization assay depicted in Figs. 4A and S2A, we used rabbit antibody recognizing the HA tag to allow us to counterstain with antibodies raised in mouse and rat.

We found that TbKifX2A-HA exhibited a strong signal further toward the proximal region of the flagellum that tapered in signal intensity in a line juxtaposed with the FAZ filament (Figs. 4B and S2D). This region of strong TbKifX2A-HA signal co-localized with the basal bodies (Figs. 4C and S2C). This pattern led us to investigate whether the kinesin was bound to the MtQ, as it originates close to the basal body and is considered a part of the FAZ (Lacomble *et al*., 2009; Sunter and Gull, 2016). To this end, we used the monoclonal antibody 1B41 recognizing the MtQ (Gallo *et al*., 1988), a somewhat contentious claim that is belied by its use as a MtQ marker in subsequent studies (Rotureau *et al*., 2011) (see Experimental Procedures for more details about the 1B41 batch used here). Indeed, TbKifX2A-HA shows co-localization with 1B41 (Figs. 4D and S2E), suggesting that TbKifX2A may associate with the MtQ.

To verify that TbKifX2A indeed binds the MtQ, we employed expansion microscopy (ExM), which enlarges cells by embedding them within a swellable polyacrylamide matrix prior to immunolabelling to allow examination of cellular structures in greater detail (Gambarotto *et al*., 2019). ExM has recently been implemented in *T. brucei* (Gorilak *et al*., 2021; Amodeo *et al*., 2021), and accentuates the part of MtQ that wraps around the flagellar pocket upon immunolabeling with antibody recognizing MT-incorporated, acetylated α-tubulin (Fig. 5A) (Gorilak *et al*., 2021). Additionally, this procedure allows the proximal MtQ to be distinguished from the nearby hook complex, which lacks MTs (Esson *et al*., 2012).

**Fig 5:**
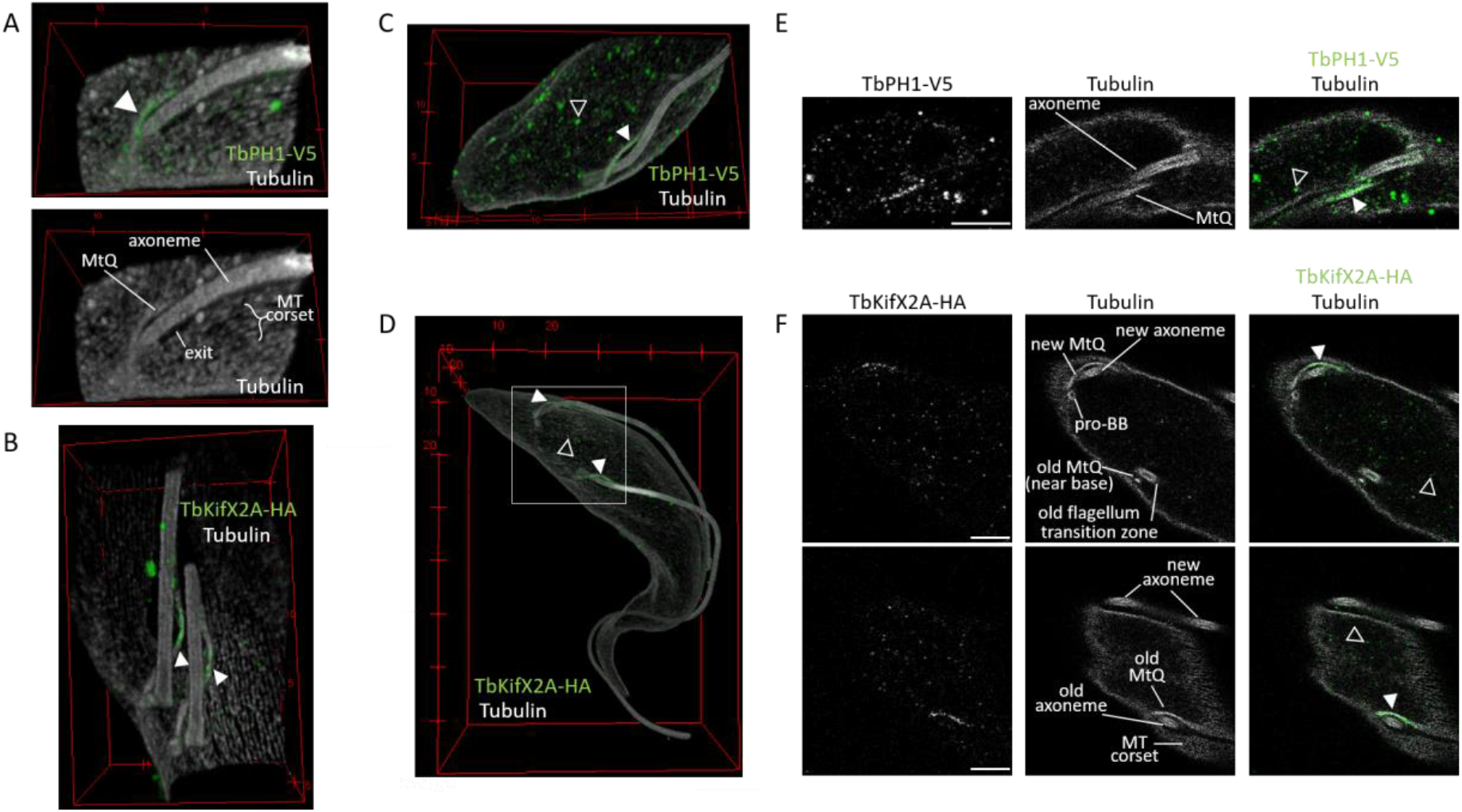
TbKifX2A and TbPH1 localize to the MtQ. Ultrastructure expansion microscopy images depicted as 3D projections (A-D) and representative confocal image sections used to make projections (E-F) of *T. brucei* expressing TbPH1-V5 and TbKifX2A-HA. (A) 3D reconstruction of the area of the cell in the vicinity of the basal body and flagellum exit from the flagellar pocket looking from outside the MT corset. Top shows signal of anti-V5 (green) and anti-acetylated α-tubulin (white) antibodies. Bottom is the same image of anti-acetylated α-tubulin signal only with relevant structures marked. Tick marks at 5 μm increments (A-C). (B) Similar reconstruction as in A but of a cell with two flagella, showing anti-HA antibody signal (green) and a view shown from inside the MT corset. (C) 3D reconstruction of the posterior part of the cell. Signal colors correspond to A. (D) 3D reconstruction of whole dividing *T. brucei* cell. Signal colors correspond to B. Box highlights region of interest shown in F. Tick marks at 10 μm increments. (E) Confocal images of a single plane from the vicinity of the basal body showing from left to right signal of anti-V5 and anti-acetylated α-tubulin antibodies(both white) plus an overlay of both signals with V5 signal in green. Left image shows 5 μm scale bar. Middle image marks relevant structures. (F) Same as in E except two confocal planes being shown, focusing on new (Top) and old MtQ (Bottom), and the left images showing anti-HA antibody signal. Solid arrowheads point to overlay of epitope antibody signal and MtQ. Hollow arrowheads point to representative epitope antibody signals found in the cytosol.

Performing ExM allowed us to localize TbPH1-V5 or TbKifX2A-HA to the part of the MtQ that spirals around the flagellar pocket, as demonstrated by the co-localization of each epitope tag with acetylated α-tubulin in both 3D reconstruction of expanded cells and individual focal planes (Figs. 5A-F). The associated V5 or HA signal was not discernible beyond the point of the MtQ joining the corset MTs, where the signal of FAZ constituents, such as ClpGM6 (Hayes *et al*., 2014), starts (Fig. S3). The fact, that the signal of TbPH1-V5 or TbKifX2A-HA was apparent only in the part of the MtQ around the flagellar pocket is consistent with this being the region of the highest concentration of both proteins in the cell based on IFA (Figs. 3D-F), as well as with a lower sensitivity of ExM due to dilution of epitopes stemming from specimen expansion. Taken altogether, ExM reveals that TbPH1 and TbKifX2A localize to the MtQ element of the *T. brucei* cytoskeleton. Additionally, we observed a punctate signal of anti-V5 or anti-HA antibody in the expanded cell body (Figs. 5C-F), consistent with the significant cytosolic fraction of both proteins (Figs. 2A, 3C-F).

### Proximity proteomics reveals the interaction network of TbPH1 and TbKifX2A

Since conventional IP of TbPH1-V5 under high salt conditions identified only TbKifX2A as its major interaction partner, we sought a different method to identify potential cargo and/or interactors of this putative kinesin complex. To do this, we tagged TbPH1 with a C-terminal biotin ligase ‘BioID2’ (Kim *et al*., 2016), appended with an HA tag (Pyrih *et al*., 2020), to facilitate proximity-dependent biotin identification (BioID) of neighbouring proteins. First, we determined by IFA on permeabilized whole cells using anti-HA antibody that TbPH1-BioID2-HA localizes adjacent to the kinetoplast, which was also observed for TbPH1-V5 (*c.f*. Figs. 2B and 6A). TbPH1-BioID2-HA also exhibited biotinylation activity, as visualized by streptavidin-conjugated to Alexa Fluor 488 (Fig. 6A), in contrast to the parental cell line lacking BioID2 (Fig. 6B). Thus, we confirmed that the chimeric protein is properly localized and appears to biotinylate TbPH1 itself or proximal proteins.

**Fig. 6:**
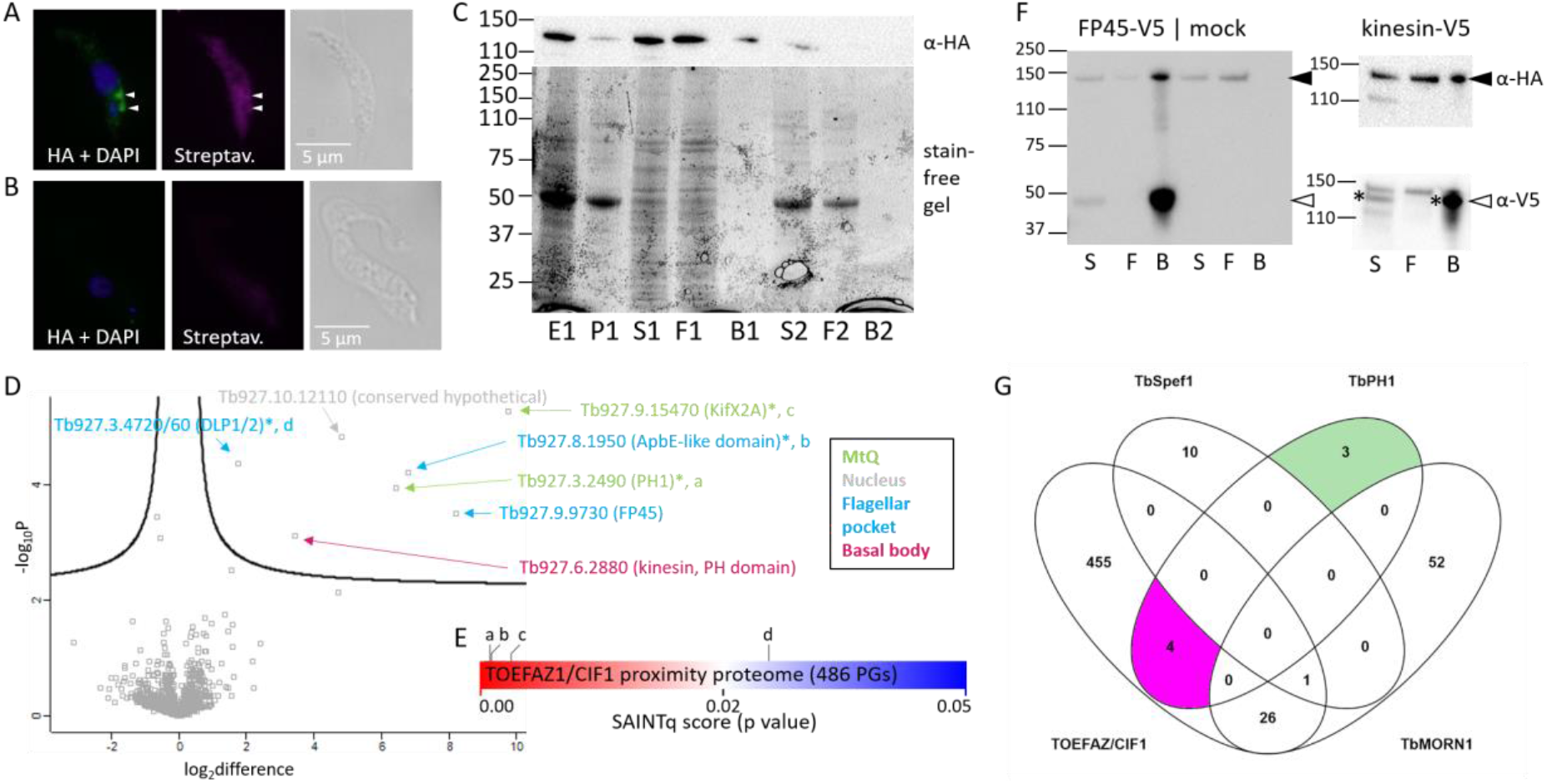
The proximal proteome of TbPH1. **(A)** IFA on whole cells showing localisation of TbPH1-BioID2-HA (green) and biotinylated proteins (magenta) and counterstained with DAPI (blue). Arrowheads denote signal in vicinity of kinetoplast, near the proximal part of the flagellum. DIC on right to show observed cell. Scale bar, 5 µm. **(B)** IFA as in **A** of parental cell line lacking TbPH1-BioID2-HA. **(C)** Western blot (top panel) and fluorescently-detected proteins resolved on SDS-PAGE as a loading control (bottom panel, stain-free gel) of a representative BioID experiment. The blot was probed with anti-HA to confirm the presence of TbPH1-BioID2-HA in the respective fractions. E: extract, P: pellet, S: supernatant, F: flow through, B: boiled beads. **(D)** Volcano plot with -10 log t-test p value plotted against the t-test difference between TbPH1-BioID2-HA sample and mock (SmOxP9 cells) from three biological replicates. Statistically significant hits are found above the curve in the top right quadrant and their identity is indicated. Accession numbers are given with additional annotations in parentheses; these are color coded based on their localization determined by TrypTag (Dean *et al*., 2017). **(E)** Heat map of SAINTq score (*i.e*. P value) of proteins interacting with TOEFAZ1 (Hilton *et al*., 2018), the relative enrichment of proteins also found in the TbPH1 proximal proteome are indicated. a:Tb927.3.2490 (TbPH1), b: Tb927.8.1950 (ApbE-like domain), c: Tb927.9.15470 (TbKifX2A), d: Tb927.3.4720/60 (DLP1/2). PG, protein groups. **(F)** Western blot showing interaction between FP45-V5/kinesin Tb927.6.2880-V5 (TbKifX2C) and TbPH1-BioID2-HA. S: supernatant, F: flow through, B: boiled beads. Black arrowhead pointing to anti-HA and white arrowhead pointing to anti-V5 signals. **(G)** Venn diagram comparing the proximity proteome of TbPH1 with those of the TOEFAZ (as depicted in **E**), the hook complex protein TbMORN1 (all 79 biotinylated proteins (Morriswood *et al*., 2013)), and the 11 high confidence proteins in the TbSpef1, a MtQ binding protein (Dong *et al*., 2020). Proteins exclusive to the TbPH1 proximity proteome are highlighted in green while magenta highlights those that are found to also be in proximity with TOEFAZ.

Purification of biotinylated proteins was performed as described elsewhere (Morriswood *et al*., 2013) (S4 Fig.). Western blotting revealed that the vast majority of TbPH1-BioID2-HA was present in the soluble S1 fraction (Fig. 6C), which was used for subsequent analysis. At first glance, this result appears to disagree with previous findings from the ‘MT sieving’ fractionation, which employs comparable amounts of non-ionic detergent and buffers of low ionic strength; however, we should emphasise that ‘MT sieving’ fractionation uses a different detergent for a shorter incubation time and at a lower temperature in comparison to the BioID purification (see Methods), which likely explains this difference.

Next, the S1 fraction was incubated with streptavidin-coated magnetic beads to capture biotinylated proteins, which were subsequently trypsinized on beads prior to liquid chromatography-tandem mass spectroscopy (LC-MS/MS). A mock control using the parental cell line lacking BioID2 expression was processed in parallel. A total of 1,131 proteins were detected with high confidence in at least one of three biological replicates (Andromeda Protein Score ≥20; >1 unique peptide per protein) (S2 dataset). Following imputation of missing values, seven proteins were shown to be preferentially biotinylated by TbPH1-BioID2-HA, as visualised by plotting the -10 log t-test p value versus the t test difference in a volcano plot (Fig. 6D). Statistically significant hits are found above the cut-off curve in this plot.

TbKifX2A was the most enriched protein based on both parameters, consistent with its strong interaction with TbPH1. As expected, TbPH1 is also among the enriched proteins, indicating that TbPH1-BioID2-HA biotinylated itself and/or a nearby untagged, endogenous form. The sub-cellular location of the five novel proteins was determined using the TrypTag protein localization resource (Dean *et al*., 2017). This allowed us to already exclude one of them (Tb927.10.12110) as a contaminant, given its nuclear localization. After TbKifX2A, a flagellar pocket protein named FP45 (Tb927.9.9730) was the next most enriched proximal protein. Interestingly, FP45 is a soluble protein that has been shown to associate with the membrane surrounding the flagellar pocket by immunogold labelling in an independent study (Gheiratmand *et al*., 2013). A putative flavin-trafficking protein (Tb927.8.1950) was the next most abundant protein, but does not have a TrypTag localization entry at this time. However, like TbPH1 and TbKifX2A, this protein was found among those that come into proximity with TOEFAZ1 (alias CIF1, (Zhou *et al*., 2018)), a protein that associates with the tip of the extending FAZ (Fig. 6E) (Hilton *et al*., 2018). Furthermore, IFA has shown that that N-terminally tagged Tb927.8.1950 exhibited a strong signal in the vicinity of the kinetoplast (Hilton *et al*., 2018). The next most enriched protein was a putative kinesin with a C-terminal PH domain (Tb927.6.2880), which is concentrated around the basal body according to TrypTag. Based on phylogenetic analysis described in the next section, we dub this kinesin TbKifX2C. Finally, two dynamin-like proteins named DLP1 (Tb927.3.4720) and DLP2 (Tb927.3.4760) were moderately enriched. DLP1/2 are localized to the flagellar pocket, which is consistent with their involvement in clathrin-mediated endocytosis (Morgan *et al*., 2004; Benz *et al*., 2017). DLP1/2 were also identified among TOEFAZ1-associated proteins, albeit with lower confidence (Fig. 6E) (Hilton *et al*., 2018).

We chose the three most enriched proteins identified by TbPH1 proximity labelling (FP45, TbKifX2C and Tb927. 8.1950) for IP experiments to verify if they represent true interactors. Cells were lysed in the same buffer as for the BioID procedure to maintain consistent conditions, except for Tb927.8.1950. Because this protein appeared to be cleaved in the BioID buffer, a different buffer that maintained the full-length protein was used for pull-down of this protein (Fig. S5). IP of the three V5 baits was successful, but TbPH1-BioID2-HA co-IP was detected only in the FP45 or TbKifX2C IPs (Fig. 6F, S5). FP45 exhibits a robust interaction with TbPH1-BioID2-HA, as the latter is noticeably depleted from the flow through fraction, which contains unbound proteins. In contrast, it appears that a fraction of TbKifX2C interacts with TbPH1, as there is a substantial amount of unbound TbPH1-BioID2-HA in the flow through. TbPH1-BioID2-HA was not detected in any of the mock IPs or the Tb927. 8.1950 IP. Thus, we conclude that FP45 and kinesin TbKifX2C are genuine TbPH1 interactors.

We also compared the TbPH1 proximity proteome to those of TbMORN1, a protein that is part of the hook complex (Morriswood *et al*., 2013), and TbSpef1, a protein localized to the MtQ (Dong *et al*., 2020) (Fig. 6G). Tellingly, there was no overlap among any of these proteomes, suggesting that TbPH1 is not in close proximity to TbMORN1 or TbSpef1 since the biotinylation radius of the BioID biotin ligase is approximately 10 nm (Kim *et al*., 2014).

### TbPH1 interacts with another X2 family kinesin in addition to TbKifX2A

Because we demonstrated that TbPH1 interacts with a kinesin other than TbKifX2A, we decided to further classify this protein, an effort that eventually led us to name it TbKifX2C. In the comprehensive survey of kinesins (Wickstead *et al*., 2010b), TbKifX2C was deemed too divergent to group into any kinesin family. Nevertheless, it does bear a canonical N-terminal, kinesin motor domain and a PH domain on its C-terminus (Fig. 1B). Furthermore, there is evidence that TbKifX2C is phosphorylated, like TbPH1 and TbKifX2A (Fig. 1A) (Urbaniak *et al*., 2012).

We noticed that the gene encoding TbKifX2C was syntenic to a *Leishmania major* gene encoding a kinesin placed in the X2 clade (Wickstead and Gull, 2006; Wickstead *et al*., 2010b). To get initial support that TbKifX2C is part of the X2 family, we performed a BLASTP search of 12 predicted proteomes covering trypanosomatid diversity (Lukeš *et al*., 2018) (Table S1A) using TbKifX2A as a query. A total of 47 protein entries were retrieved below the E-value threshold of 1e-20 (Table S1B). TbKifX2C was in this dataset along with expected hits (*e.g*. orthologs of TbKifX2A).

Encouraged by this result, we next proceeded with an unbiased molecular phylogenetic approach to test TbKifX2C’s placement within the X2 family. We retrieved from the 12 predicted proteomes 549 sequences matching the Pfam ‘kinesin motor domain’ model in a hidden Markov Model search (Table S1C, Supplemental data 1). These sequences were used to produce a trimmed alignment in which extensive gaps were removed. This alignment was then used to construct the phylogenetic tree depicted in Fig. 7A.

**Fig. 7:**
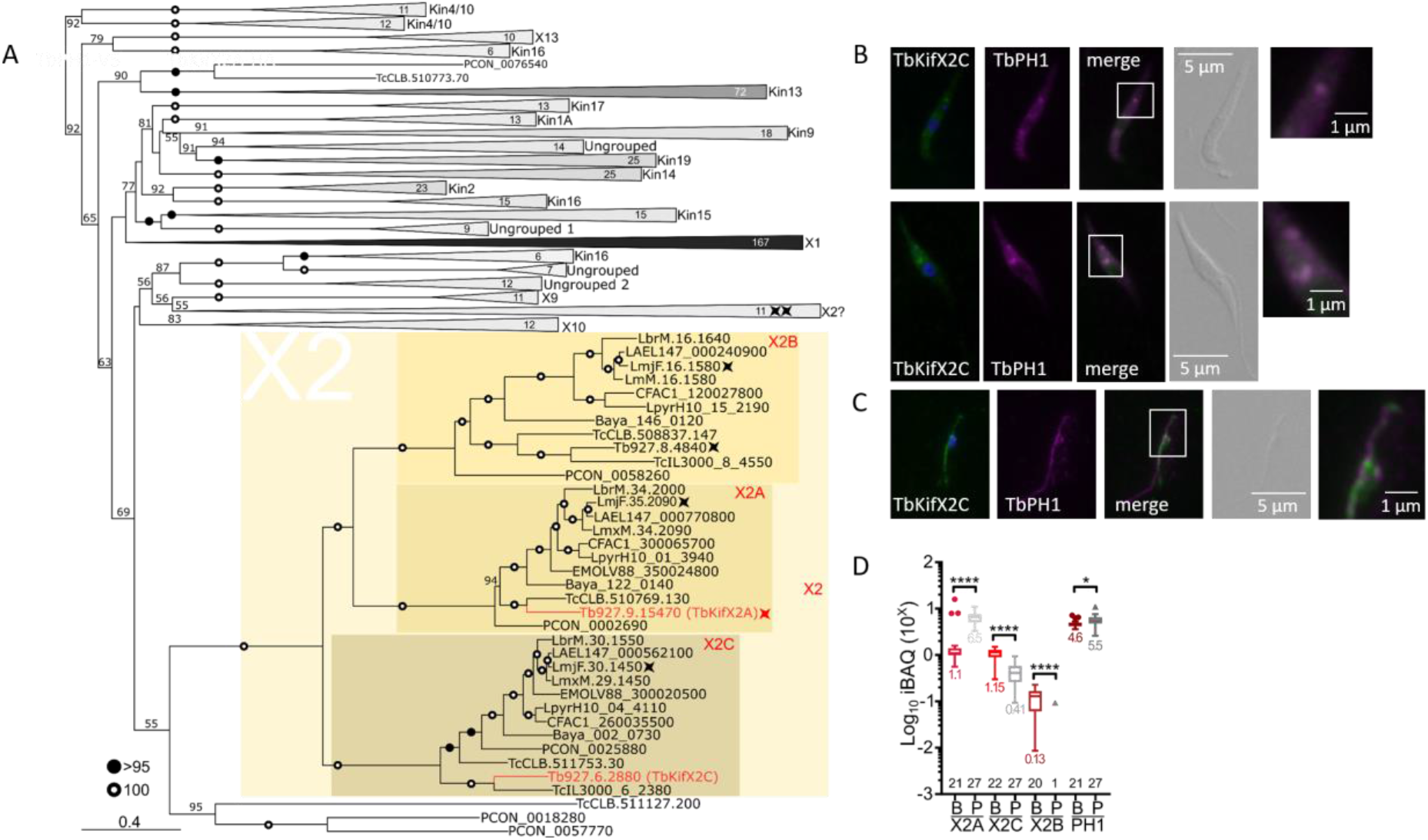
TbKifX2C is part of the X2 clade of kinesins and partially co-localizes with TbPH1. **(A)** Phylogenetic tree of 549 trypanosomatid sequences matching the ‘kinesin motor domain’ model with high confidence (see Materials and Methods). Clades containing sequences that lie outside of the X2 family were collapsed and named based on the presence of *L. major* and/or *T. brucei* representatives belonging to kinesin families as defined by Wickstead and colleagues (Wickstead and Gull, 2006; Wickstead *et al*., 2010b). The numbers of sequences belonging to each clade is shown within each triangle that is also shaded based on the number of contained sequences. Branch support values >50 are depicted; those values >95 and 100 are marked by closed and open circles, respectively. Four pointed star marks X2 family sequences as annotated by Wickstead and co-workers. Scale bar, number of substitutions per site. **(B)** IFA on whole cells (top two rows) and extracted cytoskeletons (bottom row) probing co-localisation of TbKifX2C-V5 (green) with TbPH1-BioID2-HA (magenta). Scale bar, 5 µm. Close up views of parts of strongest signal (white boxes) also shown. Scale bar, 1 μm. **(C)** Tukey plot showing differential expression of *T. brucei* X2 family kinesins in long slender bloodstream (B) and procylic (P) stages based on LC-MS/MS iBAQ values from Tinti and Ferguson (Tinti *et al*., 2022) depicted on logarithmic scale on *y*-axis. Numbers above x*-axis* report number of iBAQ spectra per sample. Colored numbers below each boxplot denote the median iBAQ value. ****, P<0.0001; *, P<0.05.

The majority of X2 family kinesins as defined by Wickstead and colleagues (Wickstead and Gull, 2006; Wickstead *et al*., 2010b) were grouped into a well-supported clade that was comprised of three distinct and well-supported sub-clades. Each sub-clade contained at least one annotated X2 family kinesin, such as TbKifX2A. Importantly, TbKifX2C was placed into the X2 clade as well, clustering with the product of the syntenic *L. major* gene. It should be noted that sequences from the basal trypanosomatid *Paratrypanosoma confusum* (Skalický *et al*., 2017) represented early diverging outgroups in all of the subclades except X2C, which contains the apparently divergent sequences from *Trypanosoma* species. As a testament to this, the *Trypansoma congolense* member of the X2C clade (TcIL3000_6_2380), which clusters with the divergent TbKifX2C, was the only X2 family sequence not retrieved in the initial BLASTP search (Table S1B). Thus, we conclude from these unbiased *in silico* approaches that like TbKifX2A, TbKifX2C is a member of the X2 family.

To see if TbKifX2C co-localizes with TbPH1, we performed IFA on fixed whole cells and extracted cytoskeletons (Fig. 7B-C). The signals from both TbKifX2C-V5 and TbPH1-BioID2-HA were enriched in the vicinity of the kinetoplast in whole cells, where they clearly overlaid (Fig. 7B). TbKifX2C IFA signal partially overlaps with TbPH1 at the part of extracted cytoskeletons near the retained kinetoplast (Fig. 7C), which is consistent with a portion of TbPH1 interacting with TbKifX2C (Fig. 6F). This partial co-localization is in contrast with the almost complete overlap of TbPH1 with TbKifX2A, which indicates that TbKifX2A is the main partner of TbPH1, perhaps explaining why TbKifX2C was not detected in the initial TbPH1 co-IP under the high salt conditions of ‘MT sieving’.

To further explain why TbKifX2C was not observed in our initial IPs identifying the TbPH1-TbKifX2A complex, we decided to look at expression of the *T. brucei* X2 family kinesins in long slender bloodstream and procyclic stages (Tinti *et al*., 2022). While TbPH1 is minimally regulated between the two life cycle stages, TbKifX2A is considerably upregulated in the procyclic stage whereas TbKifX2C is expressed at roughly a third of the levels as it is in the long slender stage (Fig. 7D). Interestingly, the *T. brucei* kinesin belonging to the X2B sub-clade (Tb927.8.4840) exhibits more regulated expression in the long slender stage, with only 1 peptide detected in the procyclic stage. Thus, it appears that TbKifX2A is the predominant X2 kinesin in the procyclic stage, which was the experimental model used in this study, which may also explain why this was the only kinesin to be bound to TbPH1 during ‘MT sieving’.

### Depletion of TbPH1 and TbKifX2A causes prominent morphological defects

To assay the function of the novel kinesin TbKifX2A and its associated kinesin-like protein TbPH1, we depleted their expression by doxycycline-induced RNAi in the cell line expressing TbPH1-V5 and TbKifX2A-HA. The downregulation of either protein alone only marginally affected the growth of *T. brucei* in comparison to the non-induced cells, despite efficient depletion of each of the respective tagged proteins (Fig. 8A, Fig. S6A). Interestingly, TbPH1 levels were also depleted in the TbKifX2A RNAi cell line (Fig. 8A, inset), suggesting that the protein might depend on its partner for stability.

**Fig. 8:**
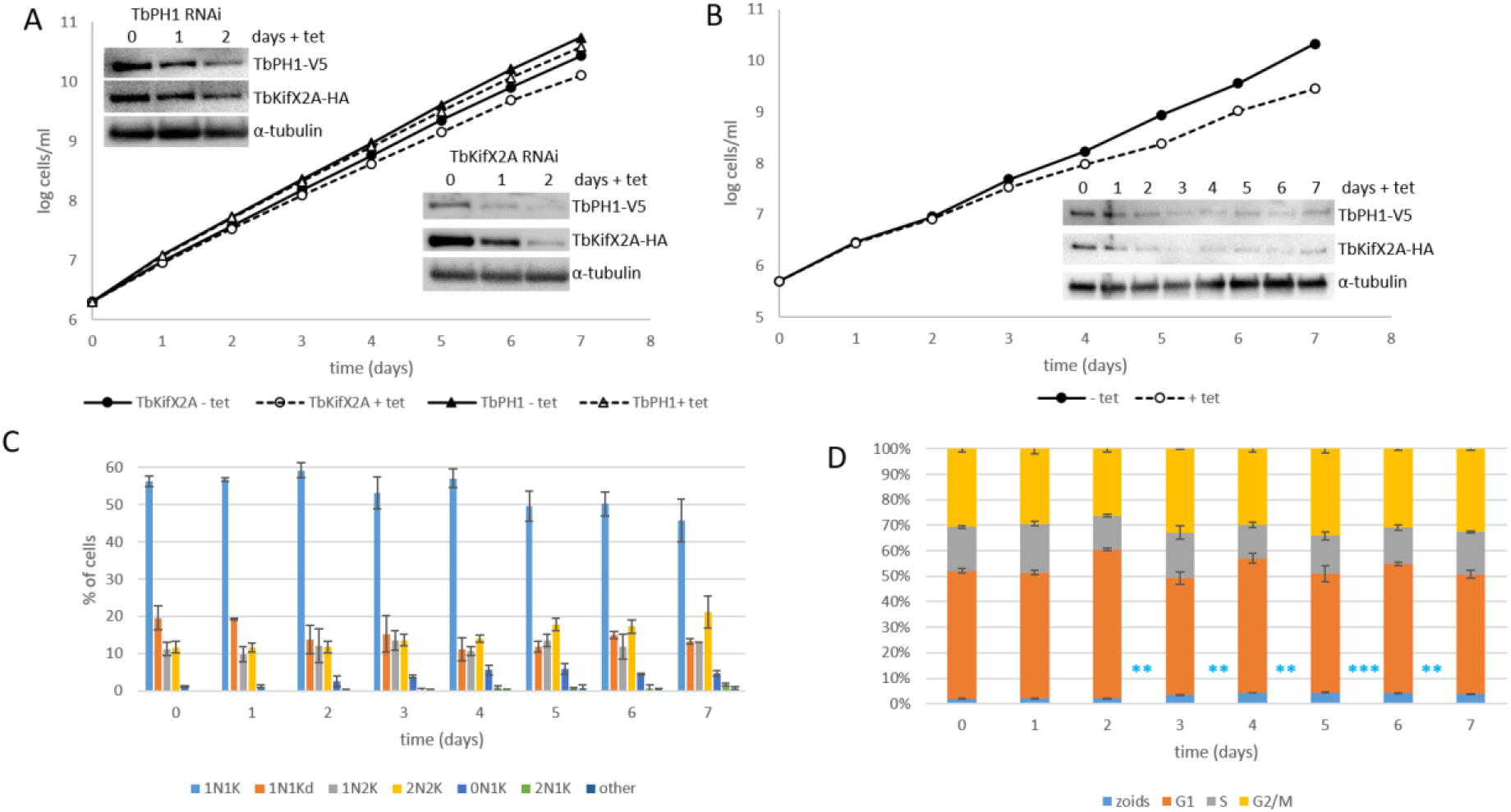
TbPH1/TbKifX2A RNAi. **(A)** Growth curves of single TbPH1 and TbKifX2A RNAi cell lines. Filled circles: TbKifX2A – tet, open circles: TbKifX2A + tet, filled triangles: TbPH1 – tet, open triangles: TbPH1 + tet. Standard deviation of measurements at each time point (n=3) is not visible at this scale. Insets show western blots probed with anti-HA (to detect TbKifX2A), anti-V5 (to detect TbPH1) and anti-α-tubulin as a loading control. **(B)** Growth curve of double TbPH1/TbKifX2A RNAi cell line. Filled circles: TbPH1/TbKifX2A – tet, open circles: TbPH1/TbKifX2A + tet. Standard deviation of measurements at each time point (n=3) is not visible at this scale. Inset depicts western blot showing depletion of TbPH1/TbKifX2A. Samples were taken at the same time as the growth curve in B was performed. The blot was probed with anti-HA (to detect TbKifX2A), anti-V5 (to detect TbPH1) and anti-α-tubulin as a loading control. **(C)** Bar chart depicting proportions of cells in different cell cycle phases during TbPH1/TbKifX2A RNAi time course. DAPI counts were performed in triplicate with at least 200 cells counted per time point. N: nucleus, K: kinetoplast. **(D)** Flow cytometry analysis of different cell cycle phases during TbPH1/TbKifX2A RNAi time course. Analyses were performed in triplicate and standard deviation is shown with error bars. Statistically significant increases in number of zoid formation compared to time point 0 is shown to the left of relevant (light blue stars; **, P<0.01; ***, P<0.001).

To get a more robust RNAi phenotype for phenotypic analysis, we generated a cell line in which both proteins were downregulated, which showed more pronounced growth inhibition than the single knock-downs (Fig. 8B, Fig. S6A). Growth slowed after 3 days of RNAi induction and did not reach levels of the uninduced counterparts at any point during the 7 day time course (Fig. 8B). Depletion of both proteins was confirmed by western blotting (Fig. 8B, inset), although neither protein was decreased below the detection limits of this assay, perhaps explaining the lack of a more severe growth phenotype.

Because both TbPH1 and TbKifX2A are modified in a way suggestive of post-translational regulation (Fig. 1A), we wondered whether the growth inhibition of the double RNAi knockdown is due to a cell cycle defect. Thus, we assayed the proportion of cell cycle stages present over a course of 7 days of TbPH1/TbKifX2A depletion, sorting individual DAPI-stained cells into different cell cycle stages by scoring their number of nuclei (N) and kinetoplasts (K) (Robinson *et al*., 1995). While we observed a moderate decrease in 1N1K cells and increase in 2N2K cells (Fig. 8C), these were not to the extent that would correspond to an unequivocal cell cycle defect. The number of zoids (cells with 1 kinetoplast and lacking a nucleus) reached a peak at day 5, making up about 7% of the population, which may suggest a cytokinesis defect. However, a conclusive cytokinesis defect phenotype should also include a concurrent emergence of 2N1K or multinucleated cells, which was not observed here (Robinson *et al*., 1995).

The cell cycle was also assayed by flow cytometry on propidium iodide-stained cells, which quantifies the portion of cells in the G1, S and G2/M phases, as well as cells with aberrant DNA content (Darzynkiewicz *et al*., 1999). While the aforementioned cell cycles phases were not markedly affected upon TbPH1/TbKifX2A depletion, this assay also revealed the statistically significant emergence of zoids from three days post-induction (Fig 8D, Fig. S6C). Thus, while we cannot conclude that TbKifX2A and TbPH1 play a direct role in cell cycle progression or cytokinesis, the two assays we employed here indicate that their depletion leads to a slight accumulation of anucleate zoids.

To gain unbiased insight into potential effects of simultaneous depletion of TbPH1 and TbKifX2A, we prepared whole cell lysates after three and five days of induction and compared them to their uninduced counterparts by LC-MS/MS. A total of 2,624 high-confidence proteins (using similar criteria as for BioID LC-MS/MS, namely an Andromeda Protein Score ≥10 and >1 unique peptide per protein) were detected for the three and five day time points, respectively. Reassuringly, TbPH1 and TbKifX2A were the most downregulated proteins at both time points as targets of RNAi-depletion (Fig. 9A). Furthermore, only two other proteins were consistently depleted at the two time points: FP45 and TbKifX2C, the two direct interactors of TbPH1. The dependence of FP45 and TbKifX2C’s stability on the presence of TbPH1 and/or TbKifX2C underscores the interaction among these proteins. Interestingly, a histidine phosphatase family protein (Tb927.11.11750) is the only significantly upregulated protein at both time points.

**Fig. 9:**
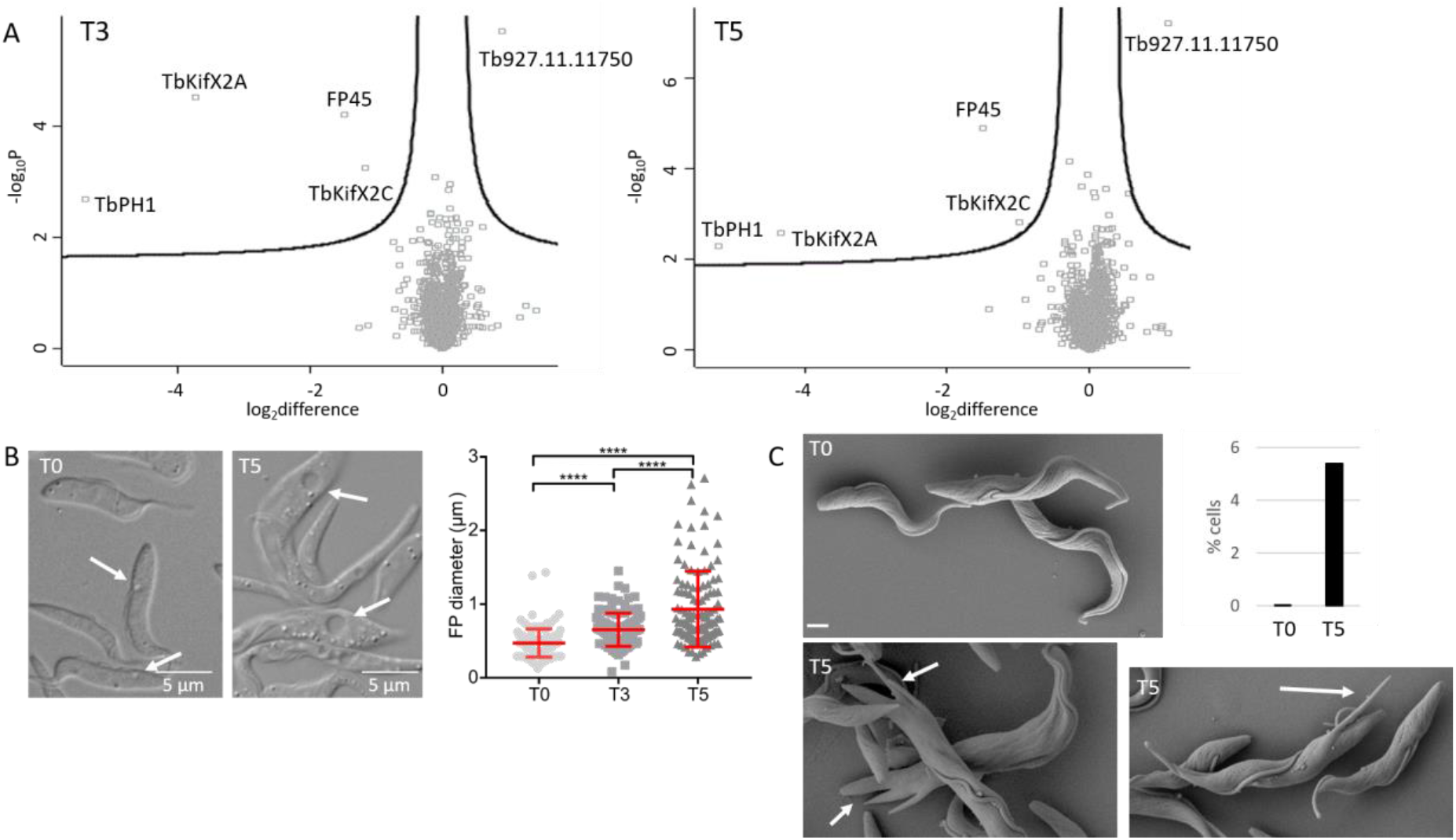
Effects of TbPH1/TbKifX2A RNAi. **(A)** Volcano plots with -10 log t-test p value plotted against the t-test difference between TbPH1/TbKifX2A RNAi sample induced for 3 (T3) and 5 (T5) days, respectively, and mock (non-induced control) from three biological replicates. Statistically significant hits are found above the curve in both top right and left quadrants and their identity is indicated. **(B)** Depletion of TbPH1/TbKifX2A causes an increase in flagellar pocket diameter. Left: Example DIC images of TbPH1/TbKifX2A RNAi cells (T0 and T5), arrows pointing at flagellar pockets. Right: Scatter plot depicting flagellar pocket diameters from 120 cells per time point. Longer middle line corresponds to the mean and whiskers are the standard deviation. Light grey circles: T0, medium grey squares: T3, dark grey triangles: T5. **(C)** Scanning electron microscopy of TbPH1/TbKifX2A RNAi cells. T0: non-induced; T5: 5 day RNAi induction. Arrows pointing to posterior end protrusions. The scale bar corresponds to 10 µm. Quantification of ‘glove’ phenotype from >380 cells per time point shown.

We next addressed whether simultaneous downregulation of TbPH1 and TbKifX2A affects cytoskeletal elements and the gross morphology of the cells. We assayed the MtQ and FAZ by IFA after five days of RNAi depletion and observed no defects in either of these structures, indicating that TbPH1 and TbKifX2A do not play a role in their biogenesis (Fig. S6B). However, by phase contrast microscopy we noticed the expansion of the flagellar pocket (Fig 9B). In contrast to the mean 472±192 nm diameter observed in 120 uninduced control cells, the flagellar pocket diameters expanded to 655±225 nm and 932±516 nm after three and five days of RNAi-induction, respectively, indicating that the severity of this phenotype increases during the RNAi-time course. In conclusion, simultaneous depletion of TbPH1 and TbKifX2A results in the expansion of the flagellar pocket, a morphological phenotype that is congruent with TbPH1 and TbKifX2A’s proximal MtQ localization.

We also noticed that at five days of RNAi-inductions, some cells exhibited protrusions extending from the posterior end. To better visualize these phenotypes, we observed these cells by scanning electron microscopy (Fig. 9C). While control cells showed normal trypomastigote morphology (Fig. 9C), we regularly observed cells with multiple protrusions (arrows) originating from the posterior end in the induced population (Fig. 9C). The emergence of this phenotype was quantified from >380 cells in SEM pictures for both induced and uninduced populations of cells. While this phenotype only occurred in about 5.4% of cells, this still represents a 10-fold increase versus non-induced cells (Fig. 9C). The occurrence of these cell types was also corroboratively found in transmission electron microscopy, which in addition provided evidence that these protrusions appear to contain normal cellular contents and were surrounded by subpellicular MTs (Fig. S7A).

This effect of TbPH1/TbKifX2A-depletion resembled the RNAi-phenotype of two tubulin-tyrosine-ligase-like (TTLL) proteins, which post-translationally modify the C-terminal tails of α- and β-tubulin by addition of polyglutamate side chains (Jentzsch *et al*., 2020). Upon depletion of either TTLL, protrusions were observed emanating from the posterior end, which prompted the authors to name this morphological change the ‘glove’ phenotype. To investigate whether TbPH1/TbKifX2A downregulation truly phenocopies that of each TTLL, we immunolabelled cells five days post RNAi-induction with antibodies recognizing glutamyl side chains (GT335 antibody). Upon depletion of either TTLL, polyglutamylation was decreased and α-tubulin tyrosination increased in the posterior end, from which the protrusions emerged (Jentzsch *et al*., 2020). We did not observe any changes in polyglutamylation upon TbPH1/TbKifX2A silencing by imaging individual cells by IFA or on the population level by western blot analysis (Fig. S7B), suggesting the mechanism underlying the emergence of posterior-end protrusions differs from that of the TTLLs.

## DISCUSSION

Kinesins and related kinesin-like proteins constitute one of the largest protein families encoded by the *T. brucei* genome (Berriman *et al*., 2005) and are represented by diverse gene repertoires in other trypanosomatids as well (El-Sayed *et al*., 2005; Ivens *et al*., 2005; Wickstead and Gull, 2006; Wickstead *et al*., 2010b). Among the eukaryotes, trypanosomatids have experienced one of the largest expansions of the kinesin motor superfamily, suggesting these proteins endow trypanosomes with unique biological properties, many of which still await elucidation (Berriman *et al*., 2005; Wickstead and Gull, 2006; Wickstead *et al*., 2010b). Indeed, this expansion has resulted in the emergence of two trypanosomatid-specific kinesin families named X1 and X2. A kinesin belonging to the former, named FCP2/TbKinX1, has been shown to maintain a connection of a new flagellum’s distal tip along the side of the old flagellum, thus facilitating the unique mechanism of templated flagellar replication in *T. brucei* (Varga *et al*., 2017). In this work, we have shed light on the thus far neglected X2 family of kinesins. First, by a widescale phylogenetic analysis using kinesin sequences from 12 trypanosomatid species, we have shown that the X2 family is comprised of 3 sub-clades. Sub-clades X2A and X2B contain kinesins from 11 of the 12 searched predicted proteomes, whereas X2C has representative from all 12. Thus, it appears that the 3 sub-clades are well-conserved in trypanosomatids, with gene losses being the exception, suggesting that this X2 kinesin repertoire is fundamental for these flagellates. Indeed, the model used in this study, *T. brucei*, possesses kinesins from all three sub-clades.

The well-supported X2 clade of our study essentially agrees with the composition of the X2 clade that was initially revealed in a phylogenetic tree of kinesins throughout eukaryotes, with one notable exception (Wickstead and Gull, 2006; Wickstead *et al*., 2010b). Two *L. major* and *T. brucei* kinesins that were originally annotated as X2 family kinesins were placed outside of the X2 clade in our analysis (marked ‘X2?’ in Fig 7A). This can be explained by the fundamental different datasets and goals of the two studies. Wickstead and colleagues aimed to understand general kinesin evolution, and thus sampled a wider range of eukaryotes, necessitating the inclusion of only two trypanosomatid species. Our study aimed to address whether a divergent kinesin should be included in the kinetoplastid-specific X2 family, and thus we restricted our dataset to a wider range of 12 trypanosomatid species. In the end, the trees are not as contradictory as they ostensibly seem at first. The outlying kinesins in our ‘X2?’ clade were placed sister to the otherwise well-supported X2 clade with less than 50% support in Wickstead’s tree (Wickstead and Gull, 2006), which is consistent with the placement of these sequences in relation to the X2 clade in our study. Thus, we propose that our tree reveals the structure of the X2 family *sensu stricto* (*s.s*.), given the level of support of this clade and contained sub-clades, implying the stability of its internal branching. The evolution of the kinesin superfamily in kinetoplastids is out of the scope of this study, but we hope this result provides impetus for future investigation of this topic.

Another property of X2 *s.s*. kinesins we have serendipitously discovered in this study is that they directly interact with TbPH1, a likely catalytically-dead, kinesin-like protein with a C-terminal PH domain that localizes to the MtQ. While we do not address here whether TbPH1 has truly lost ATP hydrolase activity, it contains two point mutations to conserved residues of the Walker A motif that are likely to ablate this activity. A glycine to proline mutation exchanges the amino acid that confers the most flexibility to the peptide backbone for one that rigidifies it. The polar threonine/serine amino acid responsible for the requisite coordination of Mg^2+^ (Deltoro *et al*., 2016) is replaced with a positively charged arginine, which would repel this cation. Divergent Walker A motifs that still retain ATP hydrolase activity do not have two such mutations at once (Hu *et al*., 2012; Deltoro *et al*., 2016). Thus, the notion that TbPH1 has an inactive kinesin motor domain is likely but empirically unproven.

We show that TbPH1 very tightly interacts with TbKifX2A in a manner persisting even at high ionic strength. This interaction most likely occurs at the part of the MtQ wrapped around the flagellar pocket, where both proteins are considerably enriched. TbPH1 and TbKifX2A exhibit a cytosolic pool as well, but it remains unknown to what extent if at all they interact away from the MtQ. Proximity labelling revealed two more direct interactors of TbPH1: FP45, a protein that localizes to the flagellar pocket (Gheiratmand *et al*., 2013) and surprisingly another X2 family *s.s*. member, TbKifX2C. Furthermore, the stability of both proteins is dependent on the presence of TbPH1 and/or TbKifX2A, indicating the interdependence of these proteins that are all enriched in the vicinity of the flagellar pocket associated MtQ.

TbPH1’s interaction with FP45 may explain the primary phenotype we have observed upon simultaneous downregulation of TbKifX2A and TbPH1: the expansion of the flagellar pocket. This phenotype is congruous with the localization of TbPH1 interactors and may be a consequence of the subsequent depletion of FP45. Unfortunately, we are not able to corroborate this result as regulatable RNAi-cell lines targeting FP45 were reported to be unstable, precluding functional analysis (Gheiratmand *et al*., 2013). Indeed, we have also encountered problems in obtaining robust phenotypes downregulating TbPH1 and TbKifX2A individually or simultaneously upon extended growth and/or cryopreservation. Thus, from our and other’s experience, complete ablation of this cohort of MtQ-associated proteins is difficult, which complicates their functional analysis.

We have also observed some morphological defects affecting a minor proportion of cells upon TbPH1/TbKifX2A downregulation that are at this time difficult to explain. One is the emergence of anucleate zoids, intensifying at the same RNAi time point when we observe the flagellar pocket expansion. At this time, we rule out that this is due to cytokinesis defects, as this would also give rise to a more conspicuous enrichment of 2N1K or multinucleate cells, which we did not observe. In the final RNAi time point examined here, we also observed cells exhibiting posterior-end protrusions, resembling the ‘glove phenotype’ observed upon depletion of two TTLL enzymes, which are responsible for post-translational modifications appending the C-terminal tails of α-and β-tubulin (Jentzsch *et al*., 2020). As neither TTLL12A (TTLL6A was not among the detected proteins, Dataset S3) nor the modifications both enzymes catalyse are affected by TbPH1/TbKifX2A-silencing, we conclude that these protrusions arise from a different mechanism.

Altogether, we conclude that TbPH1 and TbKifX2A play a still undefined role in maintaining compact flagellar pocket morphology. Whether one or both spuriously occurring defects are a direct consequence of the primary flagellar pocket expansion phenotype or due to incomplete ablation of TbPH1 and TbKifX2A remains unknown. Another possibility is that the cytosolic pool of TbPH1 and/or TbKifX2A may play a role in posterior end morphogenesis. What we can say is that it is unlikely that they are arising from secondary depletion of proteins other than the targeted ones or the downregulated TbKifX2C and FP45, as none of the other proteins detected in our LC-MS/MS analyses are affected throughout the examined RNAi time course.

TbKifX2A is the first motor protein that has been demonstrated to localize to the MtQ. This and our finding that TbKifX2C partially interacts with the MtQ-binding TbPH1 suggests the exciting possibility that the X2 family *s.s*. kinesins co-evolved with the MtQ, a cytoskeletal component that is considered to be part of the FAZ and restricted to trypanosomatids (Sunter and Gull, 2016). It has been suggested that the MtQ contains β-tubulin with a unique but still unknown modification that is recognized by the 1B41 monoclonal antibody (Gallo *et al*., 1988; Gallo and Precigout, 1988), which may at least partially explain why these kinesins and TbPH1 perferentially bind to the MtQ over the corset MTs. This specificity may be mediated by interaction with TbPH1, perhaps explaining this instance of a kinesin interacting with a catalytically-inactive kinesin-like protein. An example of this phenomenon is the budding yeast Kar3 kinesin interacting with kinesin-like Vik1, which confers to the Kar3/Vik1 complex a strong interaction with a microtubule (Allingham *et al*., 2007).

Another striking feature of the three X2 family *s.s*. kinesin paralogs in *T. brucei* is their apparent regulation in the long slender bloodstream and procyclic stages, with TbKifX2A being solely upregulated in the latter. In contrast, TbPH1 exhibits more consistent expression levels between these two stages. This suggests that the X2 *s.s*. kinesin repertoire is remodelled to adapt each stage to their specific milieus. Perhaps this regulation is due to the higher endocytic flux in long slender bloodstream forms in comparison to the procyclic forms, a process that occurs via the flagellar pocket (Field *et al*., 2009). This is certainly a worthy line of future investigation. Given that the X2 family *s.s*. kinesin may have co-evolved with the MtQ, it will be also interesting to see how the X2 kinesins have adapted to the presence of the cytostome-cytopharynx complex, an organelle that is closely associated with the MtQ in *Trypansoma cruzi* (Alcantara *et al*., 2017) and the basal trypanosomatid *P. confusum* (Skalický *et al*., 2017). Thus, our study represents a starting point for investigation of this kinetoplastid-specific family of kinesins.

## EXPERIMENTAL PROCEDURES

### Cells

Procyclic SmOxP9 cells (Poon *et al*., 2012) were grown in SDM79 with 1 µg/ml of puromycin. Cell lines bearing epitope tags were grown with 50 µg /ml hygromycin (TbPH1-V5, FP45-V5, TbKifX2C-V5, Tb927.8.1950-V5), 15 µg /ml G418 (TbKifX2A-HA) and 10 µg /ml blasticidin (TbPH1-BioID2-HA). Selection for the RNAi constructs was with 2.5 µg /ml phleomycin (TbKifX2A RNAi, TbPH1 RNAi and TbKifX2A+TbPH1 RNAi).

### Plasmids and generation of cell lines

For epitope tagging at the endogenous locus with HA and V5, long PCR products were generated with primers detailed in Table 1 using pPOT-V5-HygR and pPOT-HA-NeoR vectors as templates (based on (Kaurov *et al*., 2018) and modified from pPOTv4 (Dean *et al*., 2015)). For tagging with the BioID2 ligase (Kim *et al*., 2016), a modified version of pPOTv7-Blast-mNG (Pyrih *et al*., 2020) was used as a template for TbPH1 endogenous locus tagging. 50 µl PCR reactions were directly transfected into procyclic SmOxP9 cells using the Amaxa Nucleofactor electroporator and selection occurred approximately 16 hours after electroporation.

**Table 1.**
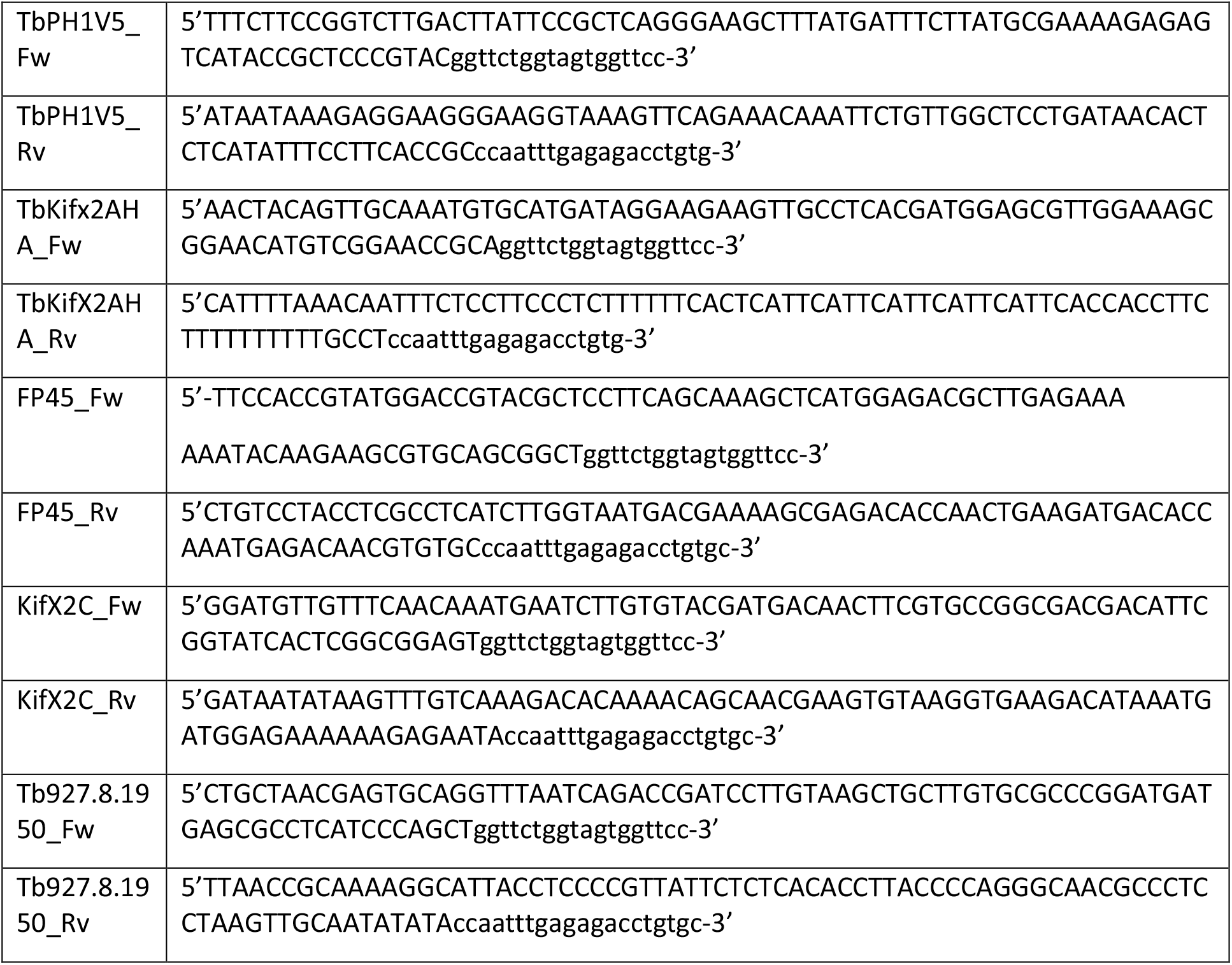

To create a vector to simultaneously downregulate both TbPH1 and TbKifX2A, a fragment of each (as suggested by RNAit2 software (https://dag.compbio.dundee.ac.uk/RNAit/) (Redmond *et al*., 2003)) was PCR amplified using primers TbPH1-RNAi-Fw (TAATtctagaGTTGCGTGATGAGCTTCAAA), TbPH1-RNAi-Rv (TATAaagcttCTCCTCTAGCACCGTTTTGC), TbKifX2A-RNAi-Fw (TAATggatccCACCGCCATTCTGCA AACAA) and TbKifX2A-RNAi-Rv (TATAtctagaGTCATTTGCGTGGGCGATAC) and cloned into BamHI-HindIII sites of p2T7-177 (Wickstead *et al*., 2002) in a three way ligation using XbaI as the restriction site in between the two RNAi fragments. The plasmid was linearized with NotI before electroporation.

### RNAi and growth curves

Procyclic form cell lines for depletion of either TbPH1 or TbKifX2A were induced at a density of 2×10^6^ cells/ml with 1 µg/ml doxycycline and growth monitored daily with a Z2 Coulter Counter Analyzer (Beckman Coulter). Cells were diluted back to the initial cell density every day. Cell lines carrying the vector for simultaneous depletion of TbPH1 and TbKifX2A were induced at a density of 5×10^5^ cells/ml with 1 µg/ml doxycycline and growth monitored daily with a Neubauer counting chamber. Cells were diluted back to the initial cell density every two days.

### DAPI counts

For cell cycle analysis, cells either induced or not for depletion of TbKifX2A and TbPH1 were spread on glass slides, the cell suspension allowed to air-dry and the slides then fixed in methanol at -20 °C. Slides were rehydrated for 5 minutes in PBS before a drop of ProLong™ Gold antifade reagent with DAPI (ThermoScientific) was added, a coverslip applied and the slide sealed. At least 200 cells were scored according to the number of nuclei and kinetoplasts per slide.

### Flow cytometry

For flow cytometry, 1-2×10^6^ cells were fixed in 30% methanol-PBS at least overnight and up to seven days at 4 °C. Samples were washed once in PBS-T (0.5% Triton) and subsequently incubated at 37 °C for 1 hour in propidium iodide staining solution (10 µg/ml propidium iodide and 9.6 µg/ml RNaseA in PBS-T, both from Sigma). Samples were analysed on a FACS Canto II (BD) collecting 50,000 events, and data processed in FCSalyzer (https://sourceforge.net/projects/fcsalyzer/).

### Cell fractionation via ‘microtubule sieving’

To entrap non-soluble and cytoskeletal proteins within the MT corset, a MT sieving technique as described in (Fritz *et al*., 2015) was employed. The experiment was conducted exactly as described before in (Fritz *et al*., 2015) (see also Fig. S1).

### Immunoprecipitation

For immunoprecipitation (IP) of TbPH1-V5 from procyclic trypanosome lysates, the V5 antibody (Thermo Fisher) was coupled and cross-linked to protein G coated dynabeads (Thermo Scientific) using 6 mg/ml dimethyl pimelidate dihydrochloride in 0.2M triethanolamine. Beads were then washed in IP wash buffer (20 mM Tris-HCl, pH 7.7, 100 mM KCl, 3 mM MgCl_2_, 0.5% TritonX-100, cOmplete™ EDTA-free protease inhibitor (Roche)) and stored at 4 °C before further usage. For proteomic analysis of TbPH1 interactions, 1×10^9^ PCF cells were pre-fractionated using the MT sieving technique described above. The SN3 lysate was incubated with V5-coupled dynabeads for 24 hours at 4 °C. Following several washes in IP wash buffer, proteins were eluted using low pH glycine buffer. Aliquots were taken of the different steps for western blot analysis. For IP of TbKifX2A-HA the same protocol was followed except that the antibody used was anti-HA (ThermoScientific) instead of anti-V5.

For IP of potential TbPH1 interactors, 2×10^8^ cells expressing FP45-V5 and TbKifX2C-V5 in addition to TbPH1-BioID2-HA were lysed and processed exactly as described in the BioID section below except that the beads used were paramagnetic V5-Trap® beads (Chromotek). Samples corresponding to 2% of the total (start and flow through fractions) and 20% of the total (beads fraction) were loaded on SDS PAGE gels, blotted onto PVDF membranes and probed with anti-HA and anti-V5 antibodies as described below.

Since full length Tb927.8.1950-V5 could not be recovered in the BioID lysis buffer, optimal lysis conditions were determined by testing different buffers (B1: 10 mM Tris-Cl, pH7.8, 150 mM NaCl, 0.1% IGEPAL; B2: 20 mM HEPES, ph 7.5, 100 mM NaCl, 0.5% CHAPS; B3: 20 mM HEPES, pH 7.5, 250 mM Nacitrate, 0.5% CHAPS). Eventually, 2×10^8^ cells expressing Tb927.8.1950-V5 in addition to TbPH1-BioID2-HA were lysed in B3 and processed as described below.

### Western blots

Cell lysates were prepared by boiling in 1x SDS sample buffer and the equivalent of 2×10^6^ cells (RNAi lines) or 4×10^6^ cells (IP samples) was loaded per well. Following western blotting, PVDF membranes were blocked for 1 hour in 5% milk-PBS. Incubation with primary antibodies (anti-V5 at 1:1,000; ThermoScientific, anti-HA at 1:1,000; ThermoScientific, anti-tubulin at 1:5,000; Sigma) was performed at 4 °C overnight. Secondary antibodies (HRP-coupled anti-mouse or –rabbit IgG; Sigma) were used at a concentration of 1:2,000 and signals were visualised using Clarity Western ECL substrate (Bio-Rad) on a ChemiDoc MP (Bio-Rad).

### SYPRO Ruby staining

Following electrophoresis, polyacrylamide gels were placed into fixing solution (7% glacial acetic acid, 50% methanol) for 30 minutes at room temperature. Following another, identical fixation step, the gel was incubated with SYPRO Ruby solution overnight at room temperature. A washing step in washing solution (7% glacial acetic acid, 10% methanol) for 30 minutes at room temperature was followed by three 5 minute washes in milliQ water before imaging on a ChemiDoc MP (Bio-Rad).

### Immunofluorescence and cytoskeleton extraction

Approximately 2×10^6^ cells were used per slide. Cells were washed in PBS and fixed in 2.3% PFA in PBS for approximately 30 minutes before being transferred to microscopy slides (ThermoScientific). Following neutralisation in 0.1M glycine in PBS, slides were incubated in methanol at -20° C overnight for permeabilisation. Slides were then rehydrated in PBS and blocked in 1% BSA in PBS for 1 hour followed by incubation with primary antibody for 1 hour in a humid chamber. The slides were then washed three times with PBS before incubation in AlexaFluor-conjugated secondary antibody (goat anti-rabbit/mouse, used at 1:1,000; Invitrogen). Following three further washes in PBS, a drop of ProLong Gold Antifade reagent with DAPI (ThermoScientific) was added, a coverslip applied and the slide sealed with nail polish. Slides were imaged with a Zeiss Axioscope or an Olympus FluoView Fv1000 confocal microscope.

For immunofluorescence analysis of cells extracted using the ‘MT sieving’ fractionation method described above the pellet fraction P2 was fixed in 2.3% PFA as described. All subsequent steps were the same as for whole cell immunofluorescence.

To extract cytoskeletons conventionally, approximately 1x 10^7^ PCF cells were collected and centrifuged at room temperature for 5 min at 800 x g and the supernatant discarded. Thereafter, the cell pellet was washed once in PBS and after resuspension in PBS applied in a drop-wise fashion to Superfrost plus® slides (ThermoScientific). The cells were left to settle before the addition of PEME buffer (100 mM Pipes, pH 6.9, 1 mM MgSO_4_, 2 mM EGTA, 0.1 mM EDTA) containing 0.5% (v/v) NP-40 (Igepal) for 10 seconds. This cytoskeleton extraction was followed by a 10 minute fixation step in 4% PFA. All subsequent steps were as described above for whole cell immunofluorescence.

#### Antibodies used in immunofluorescence

| Antibody | Dilution | Origin |
| --- | --- | --- |
| anti-HA (produced in rabbit) | 1:1,000 | Sigma, Catalog #: H6908 |
| anti-V5 (produced in mouse) | 1:1,000 | ThermoScientific, Catalog #: 377500 |
| 1B41 (MtQ)<sup>1</sup> | 1:1,000 | (Gallo *et al.*, 1988) |
| L3B2 | 1:10 | (Kohl *et al.*, 1999) |
| YL1/2 | 1:50 | (Sherwin *et al.*, 1987) |
| GT335 | 1:2000 | Adipogen Life Sciences, Catalog #: AG-20B-0020 |
| <sup>1</sup>IgM purified from hybridoma supernatant and not mouse ascitic fluid as in original report. |  |  |

### Scanning and transmission electron microscopy

For TEM, cell pellets were high-pressure frozen, freeze substituted in the presence of 2% OsO4 in acetone, and embedded into Spi-Pon812 resin (SPI) as described previously (Kaurov *et al*., 2018). For SEM of the TbPH1/TbKifX2A RNAi cell line, cells were fixed in 2.5% glutaraldehyde in 100 mM PBS overnight at 4°C and then spotted onto poly-l-lysine-coated glass cover slips. The cells were post-fixed in 2% osmium tetroxide in 100 mM PBS for 1 h at room temperature and finally washed in the same buffer. After dehydration cells were critical-point dried, coated with gold palladium and imaged.

### Expansion microscopy

Expansion microscopy samples were prepared as described previously (Gorilak *et al*., 2021). In brief, 1×10^6^ cells were fixed overnight in a solution containing 4% formaldehyde and 4% acrylamide in PBS and adhered to a poly-L-lysine (Sigma, P4707) coated coverslip. The cells were gelated for 30 min at 37°C in PBS supplemented with 19% sodium acrylate, 10% acrylamide, and 0.1% N, N’-Methylenebisacrylamide, 0.5% N, N, N’, N’-tetramethylethylenediamine, and 0.5% ammonium persulfate. The specimens were further denatured by a 2 hour incubation at 95°C in a buffer consisting of 50 mM tris(hydroxymethyl)aminomethane-hydrochloride, 200 mM sodium chloride, 200 mM sodium dodecyl sulphate, pH 9.0, and expanded by three 20 min incubations with 15 ml of ultrapure water.

Thereafter, the gels were incubated overnight first with a mixture of anti-HA (Cell Signaling Technology, 3724S; used at 1:250) and C3B9 ((Woods *et al*., 1989), used at 1:10), anti-V5 (Sigma, V8137-.2MG; used at 1:250) and C3B9 or anti-ClpGM6 ((Hayes *et al*., 2014), used at 1:200) and C3B9 antibodies, followed by washes and an overnight incubation with a mixture of secondary antibodies (Invitrogen, A11001 and A21428; used at 1:500). Both primary and secondary antibodies were diluted in PBS supplemented with 2% bovine serum albumin. Finally, the gels were washed with ultrapure water.

Prior to imaging pieces of gels containing stained expanded cells was transferred to a glass-bottom dish coated with poly-L-lysine, and imaged using a Leica TCS SP8 confocal microscope with an HC PL apochromatic 63x/NA 1.40 oil immersion objective. Z-stacks were acquired with the step size of 100 nm. Confocal z-stacks were processed in Fiji (Schindelin *et al*., 2012). 3D reconstructions were performed in the 3D viewer plugin.

### Flagellar pocket diameter measurements

For FP diameter measurements, random DIC images were taken of TbPH1/TbKifX2A RNAi cells at different time points and the diameter of 120 easily visible FPs measured in ImageJ (Rasband, W.S., ImageJ, U. S. National Institutes of Health, Bethesda, Maryland, USA, https://imagej.nih.gov/ij/, 1997-2018.).

### BioID

PCF cells (TbPH1-BioID2-HA and SmOxP9 as controls) were grown in the presence of 50 µM biotin for 24 hours. For proximity-dependent <u>bio</u>tin <u>id</u>entification (BioID), 10^9^ PCF cells were extracted in PEME buffer (100 mM Pipes, pH 6.9, 1 mM MgSO_4_, 2 mM EGTA, 0.1 mM EDTA) containing 0.5% (v/v) NP-40 (Igepal) for 15 minutes at room temperature resulting in extract E1. Following centrifugation at 3,400 x g for 2 minutes, supernatant S1 was created and pellet P1 was further processed by extraction in lysis buffer (0.4% SDS, 500 mM NaCl, 5 mM EDTA, 1 mM DTT, 50 mM Tris-HCl, pH7.4). Another centrifugation step at 16,000 x g for 10 minutes created supernatant S2. Both supernatants, S1 and S2, were then incubated with streptavidin-conjugated Dynabeads (company) for 4 hours at 4 °C. An aliquot of flow through samples F1 and F2 were retained for western blotting and the dynabeads were washed five times with PBS. A small sample of the beads was then resuspended in 2x SDS PAGE buffer and boiled, while the remainder of the beads was stored at -80 °C until further processing for mass spectrometry analysis.

### Mass spectroscopy analysis of captured biotinylated proteins

Trypsin-digestion of captured biotinylated proteins was performed on bead prior to liquid chromatography-tandem mass spectroscopy (LC-MS/MS) as previously described (Pyrih *et al*., 2020). Data was processed using MaxQuant (Cox and Mann, 2008) version 1.6.14 which incorporates the Andromeda search engine (Cox *et al*., 2011). Proteins were identified by searching a protein sequence database containing *T. brucei brucei* 927 annotated proteins (Version 51, TriTrypDB (Aslett *et al*., 2009), http://www.tritrypdb.org/) supplemented with frequently observed contaminants. Carbamidomethylation of cysteine was set as a fixed modification and oxidation of methionine and N-terminal protein acetylation were allowed as variable modifications. The experimental design included matching between runs for biological replicates. Peptides were required to be at least 7 amino acids in length, with false discovery rates (FDRs) of 0.01 calculated at the levels of peptides, proteins and modification sites based on the number of hits against the reversed sequence database. The obtained data was subsequently processed in Perseus version 1.6.14 as described in (Zoltner *et al*., 2020).

### Mass spectroscopy analysis of whole cell lysates

TbPH1/TbKifX2A RNAi cells were induced for 3 and 5 days with doxycycline, 5×10^7^ cells per replicate collected by centrifugation and washed once in 1x PBS before being snap-frozen in liquid nitrogen. Non-induced and parental control samples were also processed similarly as a control. MS LFQ analysis was performed on biological triplicates. For sample prep, SP4 (Solvent Precipitation SP3) protocol without beads was used (Harvey Johnston et al., 2021). Briefly, cell pellets (100 µg of protein) were solubilized by SDS (final concentration 1% (w/v)), reduced with TCEP [tris(2-carboxyethyl)phosphine], alkylated with MMTS (S-methyl methanethiosulfonate) and digested sequentially with Lys-C and trypsin. Samples were desalted on Empore C18 columns, dried in a speedvac and dissolved in 0.1% TFA + 2% acetonitrile. About 0.5 µg of peptide digests were separated on a 50 cm C18 column using 60 min elution gradient and analyzed in DDA mode on an Orbitrap Exploris 480 (Thermo Fisher Scientific) mass spectrometer.

Resulting raw files were analyzed in MaxQuant (v. 1.6.17.0) with label-free quantification (LFQ) algorithm MaxLFQ and match between runs feature activated (Tyanova et al., 2016). FDR was set as 0.01 at all levels. TriTrypDB-56_TbruceiTREU927_AnnotatedProteins.fasta proteome file from TriTrypDB (https://tritrypdb.org, Release 56) was used. MMTS alkylated cysteine was selected as a fixed modification (Methylthio (C), composition: H(2) C S, +45.988). Variable modifications were Oxidation (M) and Acetyl (Protein N-term). Downstream processing of the proteinGroups.txt file was performed in Perseus v. 1.6.15.0. LFQ intensity values were log2 transformed and the rows were filtered based on valid values (with a minimum of 2 in at least one group). The obtained data (comparing TbPH1/TbKifX2A cells at T3 and T5 with T0) was additionally processed in Perseus version 1.6.14 as described in (Zoltner et al., 2020).

### Bioinformatic analyses

BLASTP search was performed in TriTrypDB using KIFX2A amino acid sequence as a search query, returning hits with E=values <1e-20 . The 12 trypanosomatid predicted proteomes listed in Table S1A was searched. These 12 proteomes were chosen based on their distribution within the consensus trypanosomatid phylogenetic tree presented in Lukeš *et al*.(Lukeš *et al*., 2018).

For constructing the kinesin phylogenetic tree depicted in Fig 7A, protein sequences that match the Pfam ‘kinesin motor domain’ model (PF00225) from the same 12 predicted proteomes were mined using HMMER v3.3 (Eddy, 2009). All sequences with an E-value <1e-40 (listed in Table S1C) were taken for MAFFT alignment (Katoh *et al*., 2002). Regions of the alignment that comprised of >50% were removed. The phylogenetic tree based on the trimmed alignment was inferred with IQTree v1.3.12 with JTT+F+G4 chosen as the best fit model according to BIC (Nguyen *et al*., 2015). Branch support was estimated using ultrafast bootstrap method. Tree was visualized using FigTree software (http://tree.bio.ed.ac.uk/software/figtree/) and rooted arbitrarily.

## Data availability

The mass spectrometry proteomics data have been deposited to the ProteomeXchange Consortium via the PRIDE partner repository with the dataset identifiers PXD025802 (BioID) and PXD033717 (RNAi proteomes).

## ACKNOWLEDGEMENTS

We thank Keith Gull (University of Oxford), Linda Kohl (Muséum National d’Histoire Naturelle), Bill Wickstead (University of Nottingham), and Ziyin Li (University of Texas Medical School at Houston) for helpful comments, Susanne Kramer (University of Würzburg), Cynthia He (National University of Singapore), and Chris de Graffenried (Brown University) for antibodies, Peter Gorilak (Institute of Molecular Genetics) for assistance with imaging and image analysis and Anzhelika Butenko (Biology Centre, Czech Academy of Sciences) for advice on phylogenetic analysis. We acknowledge Karel Harant and Pavel Talacko (BIOCEV, Prague) and Zbyněk Zdráhal and David Potěšil (CEITEC, Brno) and the Proteomics Service Laboratory at the Institute of Physiology (supported by RVO, ID 67985823) and Institute of Molecular Genetics (supported by RVO, ID 68378050) of the Czech Academy of Sciences for mass spectroscopy analysis on BioID samples, excised bands and whole cell lysates, respectively. This work was supported by the Czech Science Foundation grants 20-23513S to HH, and 21-09283S and ERC CZ LL1601 to JL, the ERD funds of the Czech Ministry of Education 16_019/0000759 to HH and JL, and BioImaging grant LM2018129 to MV. Work in the VV laboratory was supported by an EMBO Installation Grant and by the J. E. Purkyne Fellowship of the Czech Academy of Sciences.

## Author contributions

Conceptualization (CB, NM, SK, VV, HH), formal analysis (CB, NM, SK, HV, MV, VV, HH), investigation (CB, NM, SK, HV, MV), visualization (CB, VV, HH), writing – original draft (CB, HV, JL, VV, HH), funding acquisition (JL, VV, HH), project administration (HH).

## FIGURE LEGENDS

**S1 Fig.:**
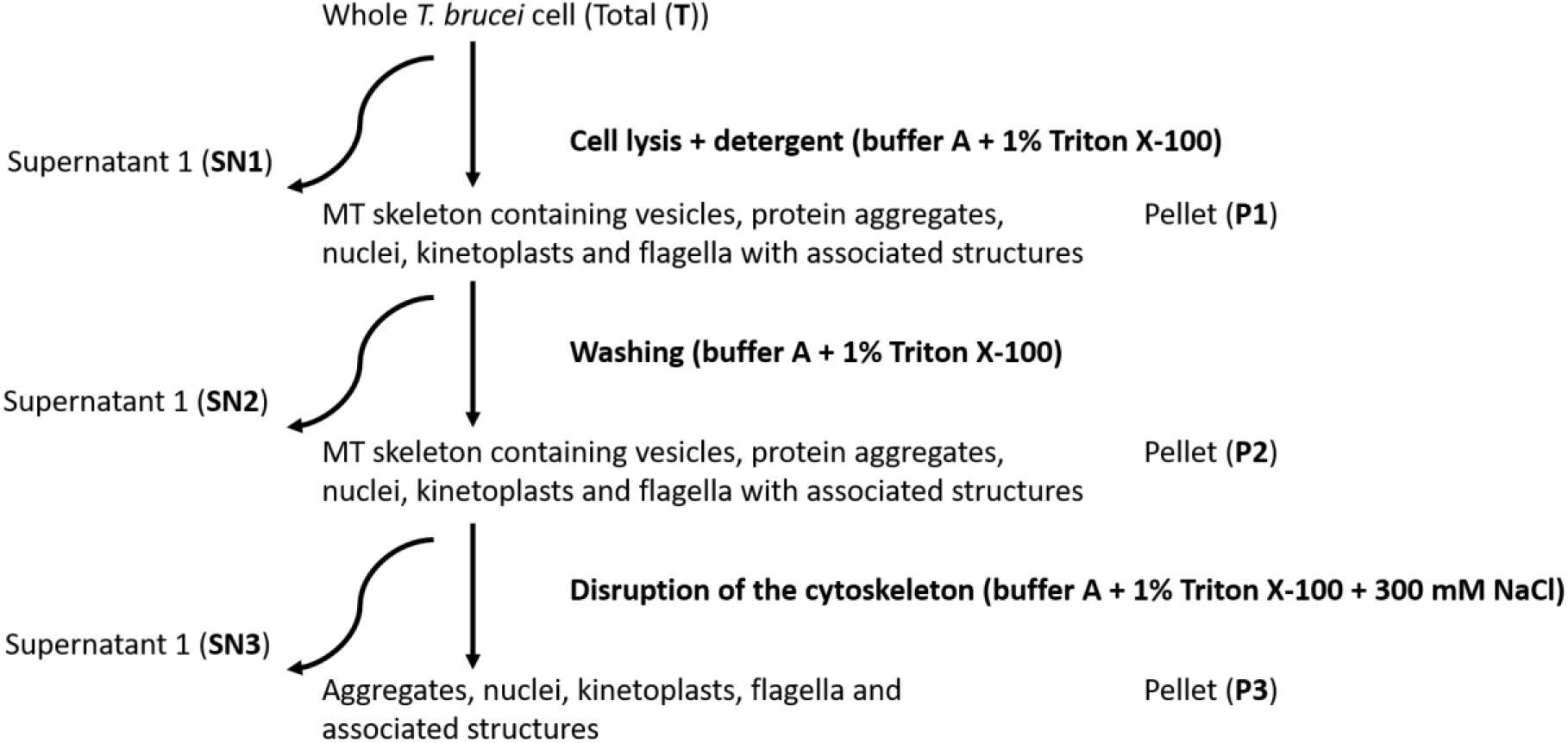
Schematic depiction of microtubule sieving technique. According to (Fritz *et al*., 2015).

**S2 Fig.:**
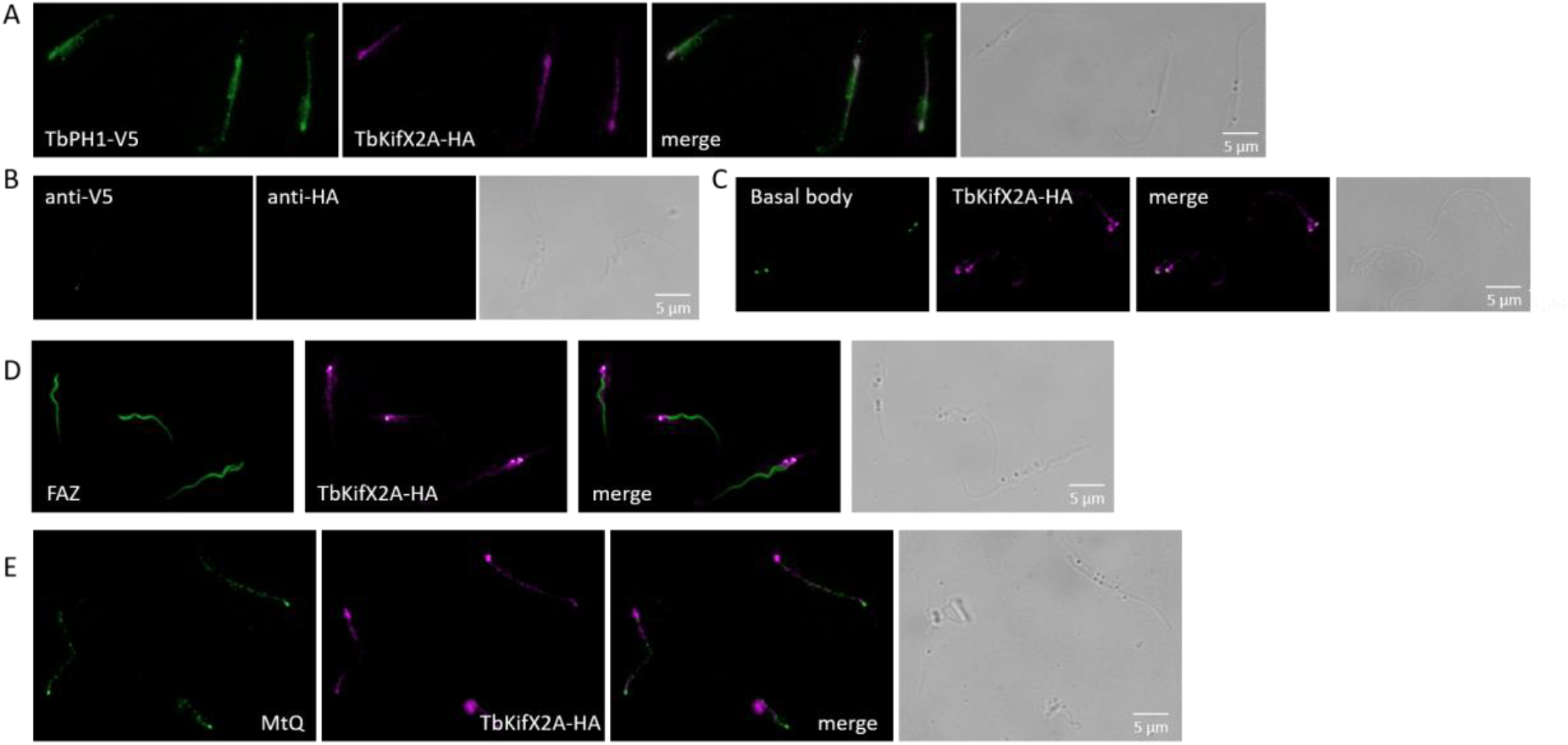
Wider field of view of multiple extracted cytoskeletons co-localizing TbKifX2A and TbPH1 to basal body **(C)**, FAZ **(D)** and MtQ **(E)**. Images depicted as in Fig 4 with scale bar corresponding to 5 µm. **(A)** Co-localization (merge, white) of TbPH1-V5 (green) and TbKifX2A-HA (magenta). **(B)** Parental cell line lacking TbPH1-V5 and TbKifX2A-HA, incubated with antibodies recognizing V5 and HA epitopes used to acquire other IFA images.

**S3 Fig.:**
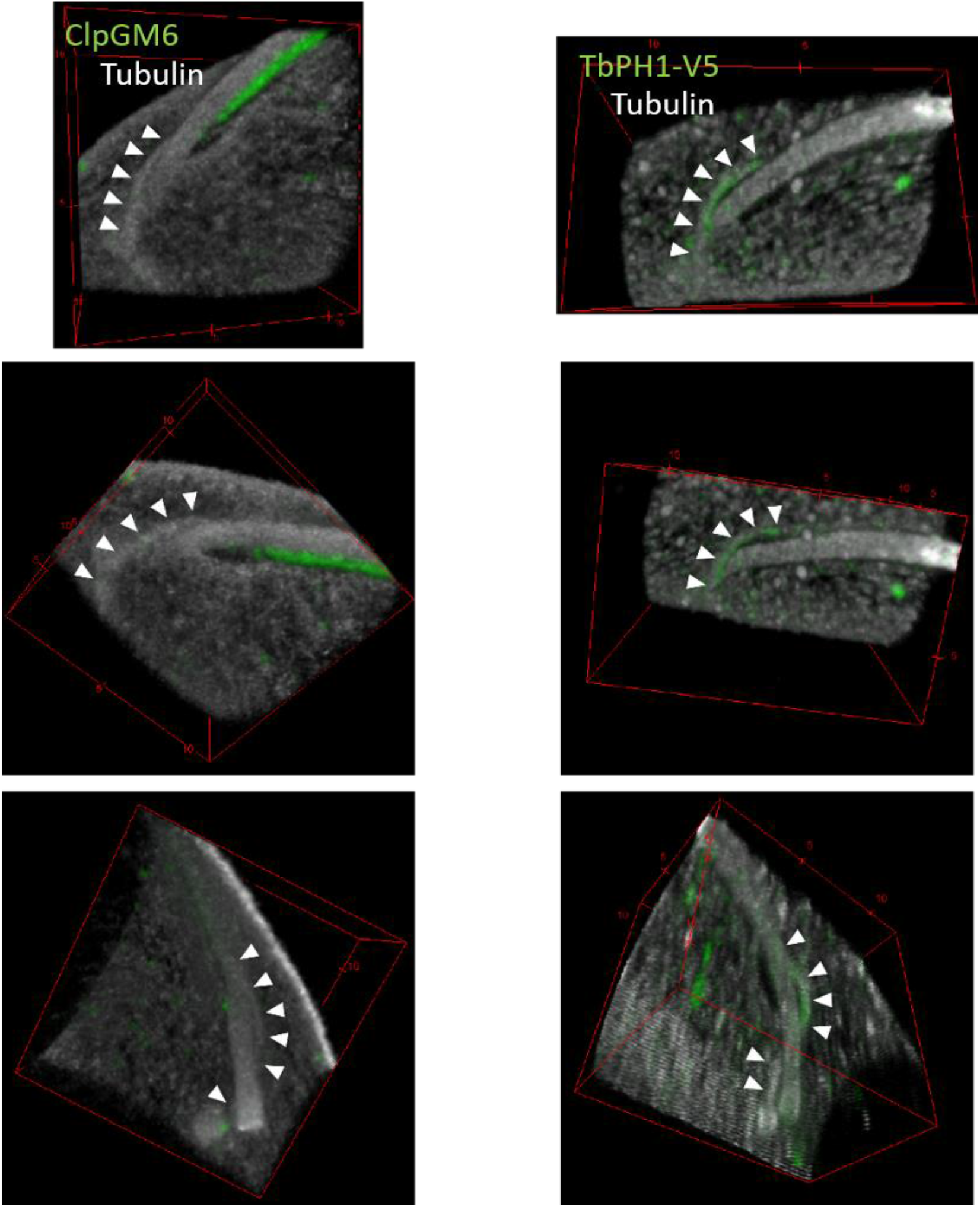
Ultrastructure expansion microscopy of a cell labelled with anti-ClpGM6 (to detect the FAZ, green) and anti-acetylated α tubulin antibody C3B9 (grey). Only the region around the flagellar pocket and flagellum exit shown. Similar region of a cell stained for TbPH1-V5 from Fig. 5A shown for comparison. The areas are shown under different angles. Arrowheads indicate the part of the MtQ inside the corset.

**S4 Fig.:**
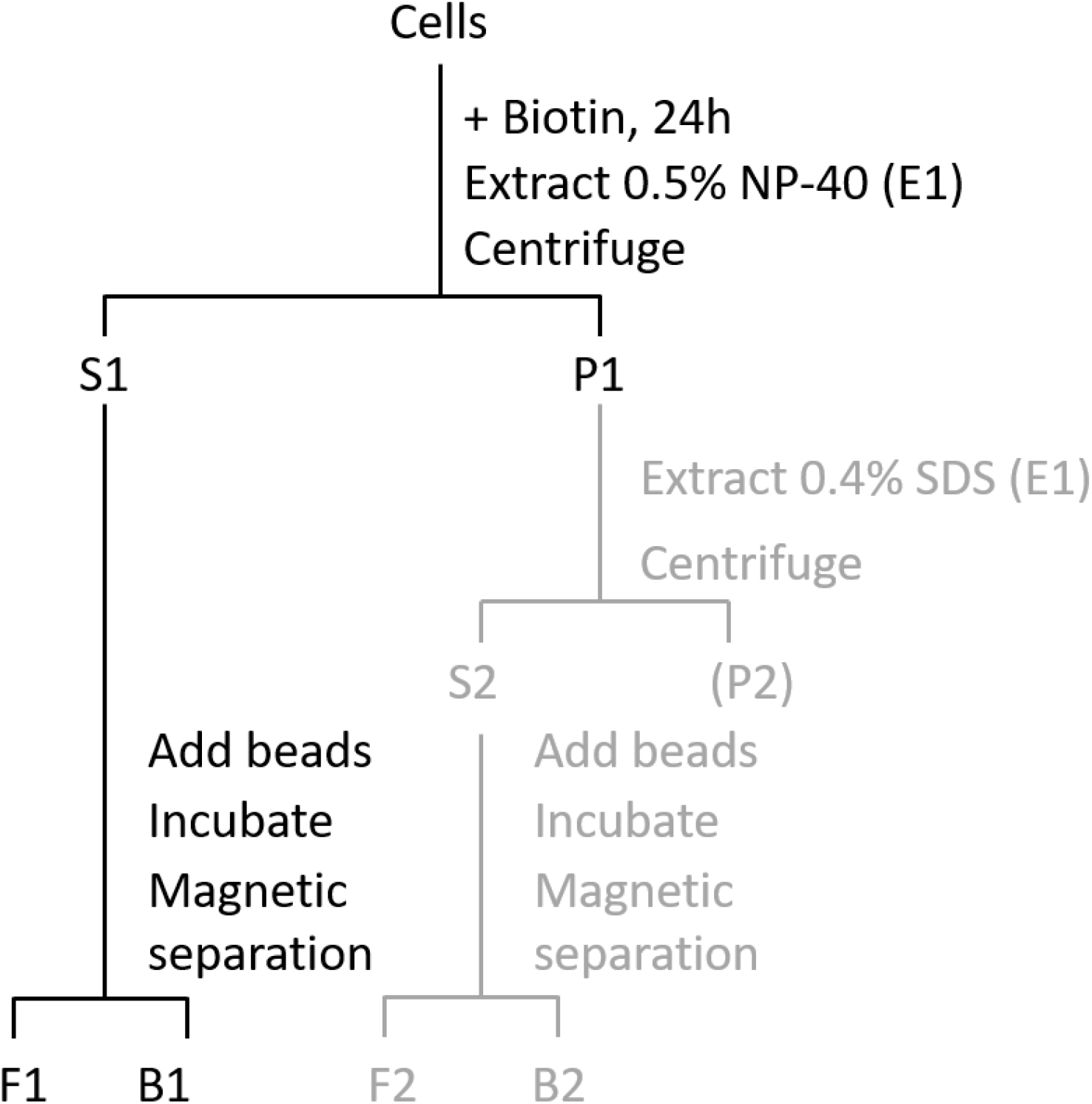
Schematic depiction of proximity-dependent biotinylation (BioID). According to (Morriswood *et al*., 2013). Only steps in black were performed in our study.

**S5 Fig.:**
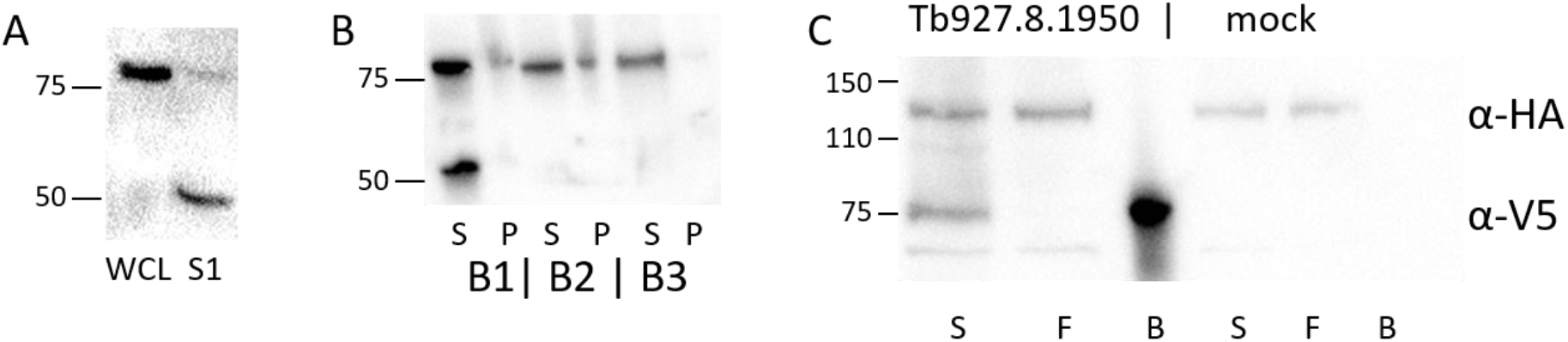
Tb927.8.1950-V5. **(A)** Western blot showing size difference between Tb927.8.1950-V5 in whole cell lysates (WCL) and BioID S1 fraction (S1). **(B)** Western blot showing size and solubility in different lysis buffers. S: soluble, P: pellet, B1: 10 mM Tris-Cl, pH7.8, 150 mM NaCl, 0.1% IGEPAL; B2: 20 mM HEPES, ph 7.5, 100 mM NaCl, 0.5% CHAPS; B3: 20 mM HEPES, pH 7.5, 250 mM Nacitrate, 0.5% CHAPS. **(C)** Western blot showing IP of Tb927.8.1950-V5. The blot was also probed for the presence of TbPH1-BioID-HA. S: supernatant, F: flow through, B: boiled beads.

**S6 Fig.:**
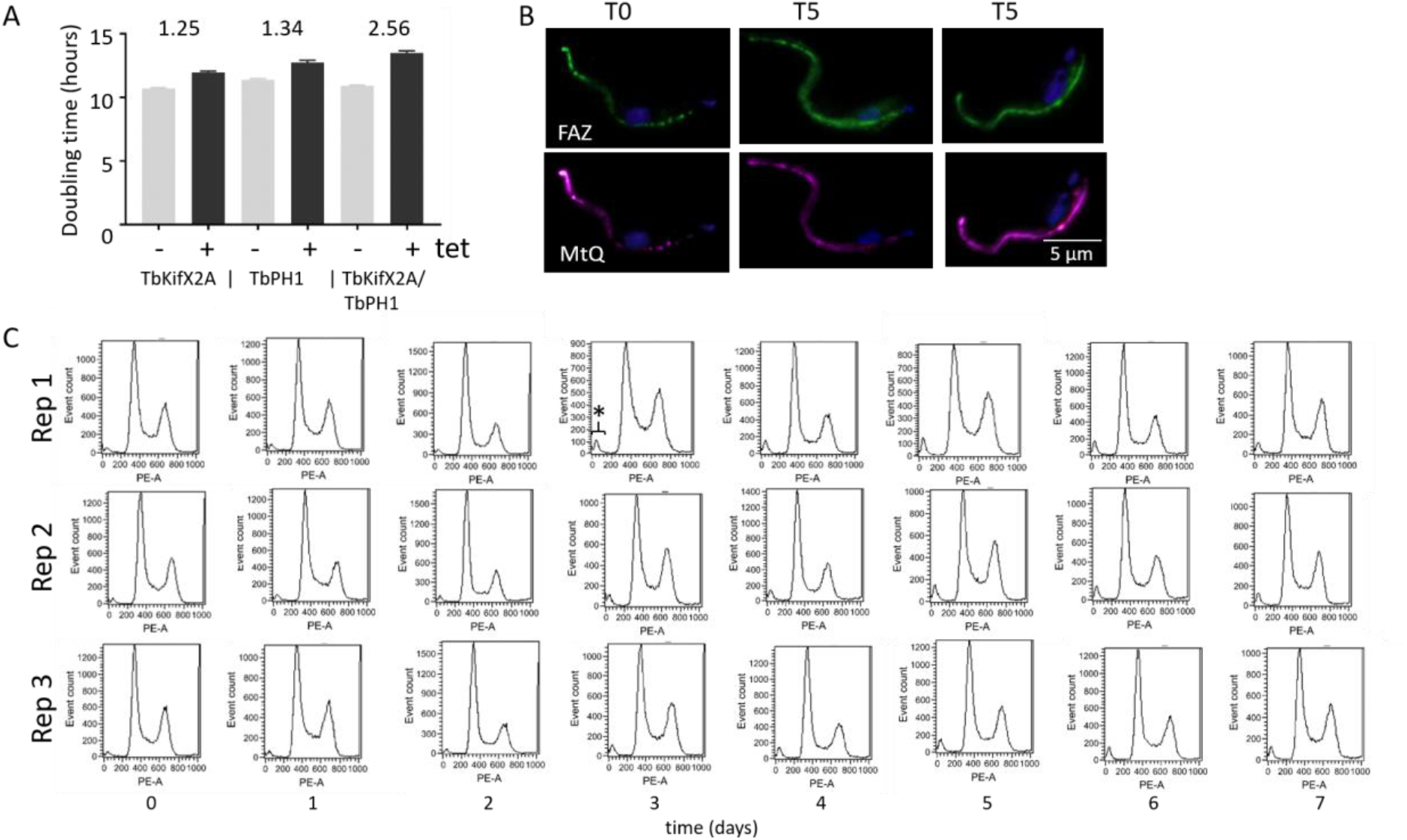
**(A)**Bar chart showing population doubling times for TbPH1, KifX2A and TbPH1/KifX2A depletion cell lines. -/+: -/+ doxycycline to induce RNAi, the average difference in hours from three experiments is shown above the respective bars. Standard deviation shown using whiskers. **(B)** IFA of TbPH1/TbKifX2A RNAi cells. T0: non-induced, T5: induced for 5 days. Extracted cytoskeletons were labelled with L3B2 (recognising the FAZ filament, green), mAB 1B41 (recognising the MtQ, magenta) and DAPI (blue). Scale bar, 5 µm. (C) Flow cytometry histograms of TbPH1/TbKifX2A RNAi cells. Three replicates are shown, asterisk marks the appearing zoid peak in one example histogram.

**S7 Fig.:**
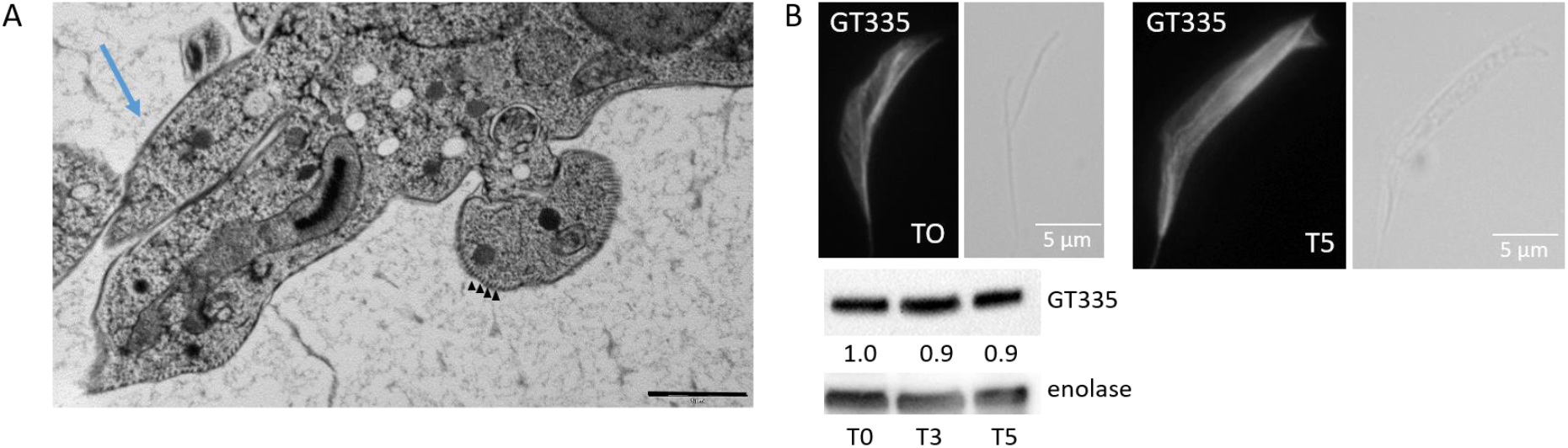
Transmission electron microscopy and IFA of TbPH1/TbKifX2A RNAi cell line. Cells were induced for 5 days. **(A)** TEM image depicting posterior end protrusions (arrow). Scale bar, 1 µm. **(B)** Top: IFA on cytoskeletons probing effects of TbPH1/TbKifX2A depletion on post-translational modifications of tubulin. Cells were stained with GT335 (white, staining glutamyl side chains). The scale bar corresponds to 5 µm. Bottom: Western blot of cells taken after 3 to 5 days of growth in presence (T3, T5) or absence (T0) of doxycycline probed with antibodies recognizing GT335 and α-enolase (Hannaert *et al*., 2003), the latter used as a loading control.

**Table S1:** Bioinformatic analysis of X2 family kinesins. **(A)** List of predicted trypanosomatid proteomes used in this study with the protein name prefixes used to name utilized sequences. **(B)** List of proteins retuned by BLASTP with E-value<1e-20 by searching the predicted trypanosomatid proteomes with TbKifX2A query sequence. **(C)** List of 549 trypansomatid sequences that match the ‘kinesin motor domain’ model (Pfam PF00225) with E-vale<1e-40 by HMMER search (Eddy, 2009).

**S1 Dataset:** Mass spectrometry analysis of excised bands B1 and B2 from Fig. 3B.

**S2 Dataset:** Mass spectrometry data of TbPH1 BioID experiment. Data with and without imputation and filtered for Andromeda scores >20 and presence of >1 unique peptide, which was used to generate the volcano plot in Fig. 6D.

**S3 Dataset:** Mass spectrometry data of whole cell lysate TbPH1/TbKifX2A RNAi experiment. Data filtered for Andromeda scores >10 and presence of >1 unique peptide, which was used to generate the volcano plots in Fig. 9A.

